# Nuclear basket localized proteasomes maintain circadian period through nuclear TOC1 proteolysis

**DOI:** 10.64898/2026.05.21.727017

**Authors:** Yeon Jeong Kim, Breagha Magill, Jing-Wen Yao, Hua Shi, Yun Sun Lee, Iris Meier, David E Somers

**Affiliations:** Department of Molecular Genetics, The Ohio State University, Columbus OH 43210 USA; State Key Laboratory for Conservation and Utilization of Bio-Resources in Yunnan, Yunnan Agricultural University, Kunming, 650201, China; Crop Biotechnology Institute, Institutes of Green-bio Science and Technology and Department of International Agricultural Technology Seoul National University, Pyeongchang, Republic of Korea

## Abstract

Post-translational control of circadian period can involve changes in protein intracellular localization to affect clock function. Many clock proteins rely on a nuclear presence for their activity. As the primary gateway regulating the movement of molecules between the cytosol and the nucleus, the nuclear pore may assist in circadian system maintenance. We describe roles for the nuclear basket proteins, NUA and NUP136, in the maintenance of *Arabidopsis* circadian period, through effects on the stability of the core clock protein, TOC1. The circadian period of *nua* and *nup136* mutants is significantly longer than that of wildtype plants. We show that NUP136 interacts with NUA, proteasome components and TOC1 *in vivo,* recruiting them to the inner nuclear rim. TOC1 interaction with the NUP136-NUA complex leads to a proteasome-dependent degradation of TOC1. Loss of *NUP136* or *NUA* disrupts this regulatory environment, leading to aberrant nuclear TOC1 accumulation and consequent lengthening of circadian period. Our work thus identifies nuclear basket-localized proteasomes as key to the maintenance of circadian period.

## Introduction

The circadian clock is an intrinsic timekeeping mechanism that enables organisms to sustain their cellular processes with synchronization to diurnal cycles of light and temperature. The approximately 24 h rhythm administers physiological events in an anticipative and buffered manner helping to confer adaptive advantage and enhanced fitness to organisms under changes in daily environmental cues ^1,2^. The benefit likely arises from discrete phasing of output gene expression, which allows the appropriate timing of metabolic and developmental processes.

In *Arabidopsis thaliana*, the fine-tuning of clock pathways is controlled by multiple layers of interlocked feedback loops^3^. Transcriptional regulatory circuits are relatively well defined, which were first characterized as mutually inhibitory interactions between transcriptional repressors, two partially redundant MYB domain-containing transcription factors, *CIRCADIAN CLOCK ASSOCIATED 1* (*CCA1*) and *LATE ELONGATED HYPOCOTYL* (*LHY*), and a pseudoresponse regulator (PRR), *TIMING OF CAB EXPRESSION1* (*PRR1/TOC1*) ^4–7^. Other *PRR* family members, *PRR9*, *PRR7* and *PRR5* sequentially express throughout the day and participate in the central loop to restrain the expression of *CCA1* and *LHY* to near dawn ^8–11^. De-repression of *CCA1* and *LHY* are mediated by evening complex (EC) genes, *EARLY FLOWERING3* (*ELF3*), *ELF4*, and *LUX ARRHYTHMO* (*LUX*) that down-regulate the expression of *PRR9* and *PRR7* at night^12–15^.

Recently, additional transcriptional feedback loops and post-transcriptional/translational processes have been added to the core transcriptional network^16^, including molecular mechanisms which underlie the nuclear translocation of particular clock proteins^17,18^. For example, PRR5 enhances the nuclear abundance of TOC1 through binding to the TOC1 N-terminus^19^. The nuclear localization of ELF3 and the sequestration of GIGANTEA (GI) to nuclear bodies are both promoted by ELF4, which elevates ELF3 nuclear protein levels^14^ and controls subnuclear sequestration of GI nuclear foci^20^. These examples demonstrate that the formation of clock protein complexes is important for their nuclear presence, suggesting these interactions enhance or facilitate nuclear import or alters nuclear protein proteolysis. Additionally, many nuclear-localized clock proteins possess nuclear localization signals (NLS) recognized by nuclear import receptors (karyopherins) which regulate their nuclear import through the nuclear pore complex ^21,22^.

The nuclear pore complex (NPC) is a large assembly of more than 30 evolutionarily conserved proteins (nucleoporins, nups) that span the nuclear envelope, connecting the cytoplasm and the nucleoplasm ^23,24^. It consists of a number of subcomplexes, organized into central inner and outer pore rings, cytoplasmic filaments, and a nuclear basket which extends into the nucleoplasm ^25,26^. The center of NPC is filled with phenylalanine–glycine (FG) repeats-containing Nups, and its meshwork nature enables the NPC to act as a permeability barrier. Various functions and interactions of the Nups have been proposed through cryogenic electron microscopic and tomographic structural analysis (cryo-EM/ET), NPC reconstructions and mutant analyses. The nuclear pore and basket have versatile roles beyond the regulation of nucleocytoplasmic shuttling ^25,27–30^.

In Arabidopsis, the nuclear basket protein *NUCLEAR PORE ANCHOR* (*NUA*) has been implicated in flowering time control and auxin signaling ^31,32^. *nua* mutations are impaired in mRNA nuclear export and show enhanced levels of SUMO conjugates, features shared with other nuclear pore-associated mutants ^33–35^. Here, we have examined the molecular basis of the circadian period defect of *nua* mutants, which display long free-running periods. We identify an aberrantly high nuclear level of the transcriptional repressor TOC1 as the primary cause for period lengthening and identify reduced TOC1 nuclear proteolysis as the basis for this accumulation. We establish that a NUA-NUP136 complex, that comprises a principal component of the basket array, can recruit proteasomes to the nuclear rim and propose the absence of either basket protein diminishes the basket-associated proteasome population, allowing an increase in TOC1 protein and a longer period. Hence, we establish how nuclear basket-localized proteasomes are essential to the maintenance of circadian period in Arabidopsis.

## Results

### *nua* mutants lengthen circadian period through aberrant TOC1 accumulation

Previous work reported that mutations in the nuclear basket protein, NUA/AtTPR, show pleiotropic effects, including early flowering ^31,32^. As early flowering can be indicative of defects in the circadian system, we examined the free running period of *nua-3* and *nua-4* alleles. Using clock promoter-luciferase reporters (*CCA1::LUC*; *PRR7::LUC*), we found the periods of both alleles are approximately 1.5 to 1.7 h longer than wildtype (CCA1::LUC: WT: 24.0 h; *nua-3*: 25.4 h; *nua-4*: 25.7 h), with the null allele (*nua-4*) slightly more severe (Fig. 1A). qPCR analysis of the phasing of select clock genes confirmed this long period finding (Fig.1B).

**Figure 1.**
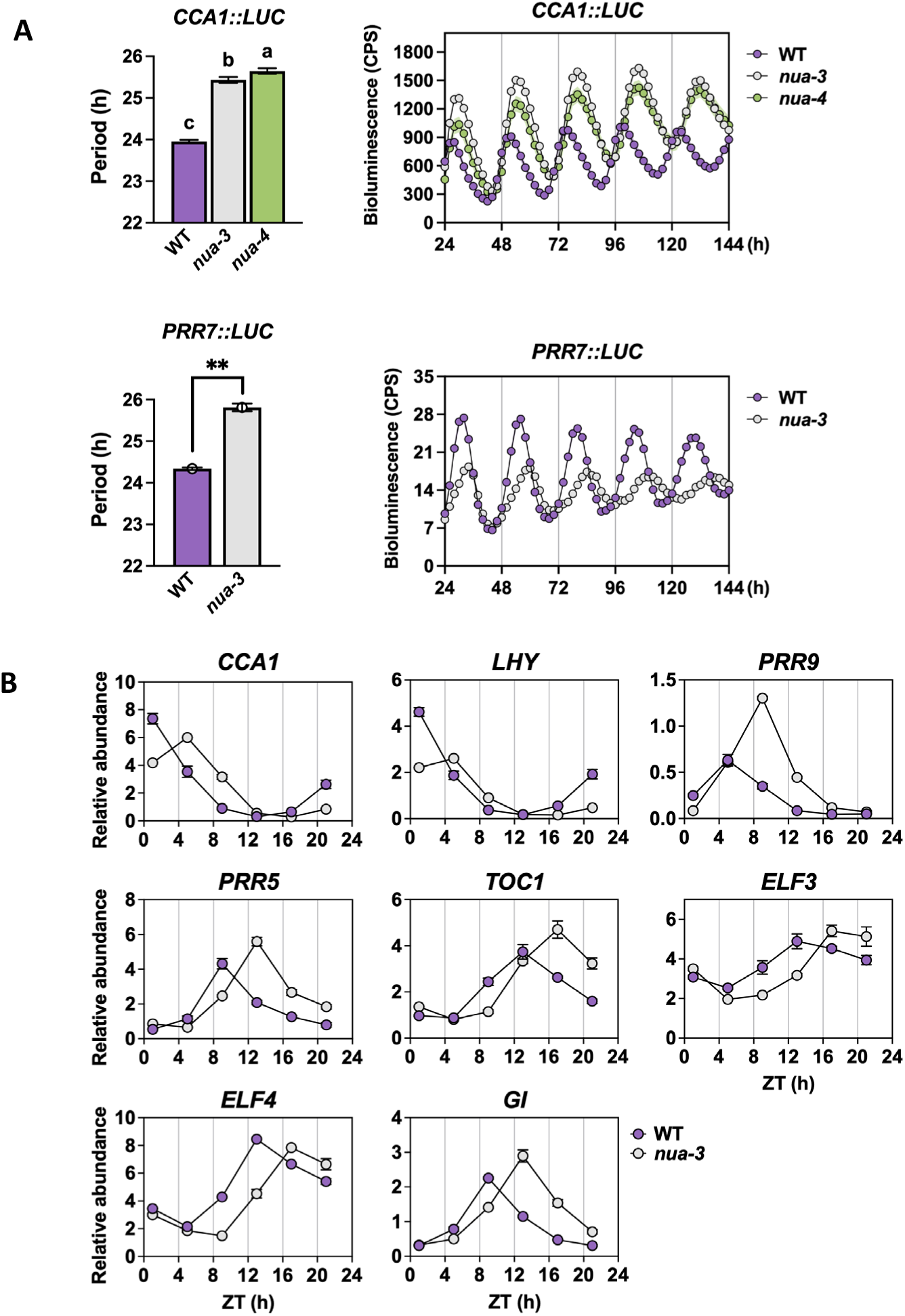
*nua* mutants lengthen circadian period. **(A)** Bioluminescence assays of *CCA1::LUC* (top) and *PRR7::LUC* (bottom) reporter lines in wild type (WT), *nua-3*, and *nua-4* backgrounds under continuous light conditions. Period estimates are shown as bar graphs. A representative result from three biological trials is shown (*CCA1::LUC*, n ≥ 15; mean ± SD) or from two biological trials (*PRR7::LUC*, n ≥ 16; mean ± SD). Asterisks indicate statistically significant differences compared to WT (\*\**P* < 0.01, Student’s t-test). **(B)** Quantitative RT-PCR analysis of circadian clock gene expression in WT and *nua-3* seedlings under continuous light conditions. Transcript levels of *CCA1*, *LHY*, *PRR9*, *PRR5*, *TOC1*, *ELF3*, *ELF4*, and *GI* were measured at indicated zeitgeber time (ZT) points and normalized to *UBQ*. Purple circles, WT; gray circles, *nua-3*. Data represent one biological trial. Values are shown as relative abundance.

*nua* mutants can be characterized, in part, by an aberrantly high nuclear level of SUMO conjugates ^31,32^. To test whether the long circadian period arises from altered SUMO conjugation we tested the *nua-3 siz1-2* double mutant, which eliminates nuclear conjugate accumulation (Fig. 2A). SIZ1 is an E3 SUMO ligase which has been shown to affect circadian period in plants ^36^, and we have confirmed a slight period shortening in *siz1-2* (Fig. 2B). If altered SUMO conjugation was responsible for the long period of *nua* mutants, the *siz1* circadian phenotype (23.8 hrs) should be epistatic to the long *nua* period (25.8 hrs). Instead, the *nua-3 siz1-2* period (25.1 hrs) is close to *nua* single mutant (Fig. 2B), indicating that the SUMOylation status of the cell is not the primary cause of the long period in *nua* backgrounds.

**Figure 2.**
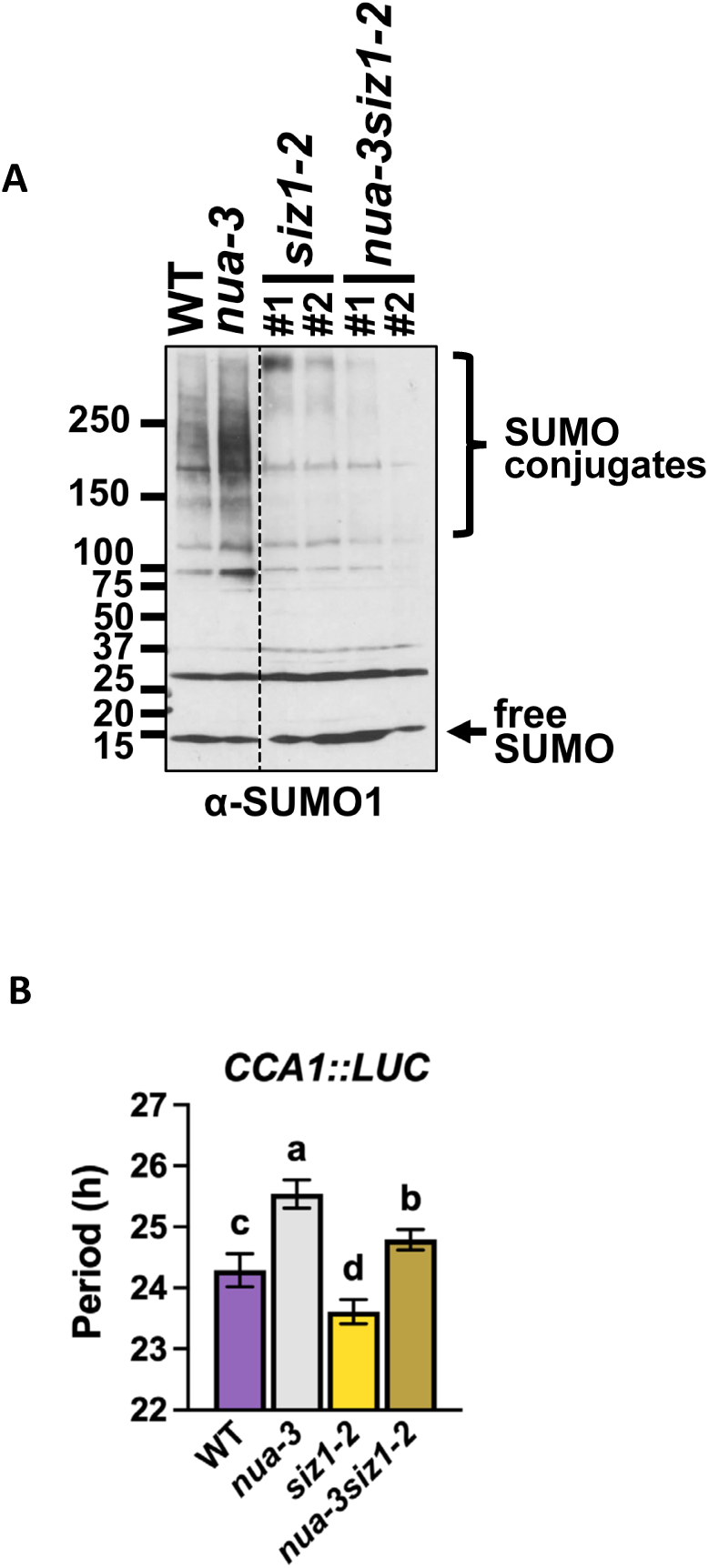
Altered SUMO conjugation levels are not primary cause of long period in *nua* mutants. **(A)** Immunoblot analysis of SUMO conjugation in WT, *nua-3, siz1-2,* and *nua-3siz1-2* mutant seedlings. Total protein extracts were immunoblotted with an anti-SUMO1 antibody. The bracket denotes high-molecular-weight SUMO conjugates. Two independent segregants were tested for *siz1* and *nua-3 siz1*. **(B)** Quantification of *CCA1::LUC* bioluminescence period length in the indicated genotypes. A representative result from two biological trials is shown (*n* ≥ 10; mean ± SD). Different letters indicate statistically significant differences (*P* < 0.05, one-way ANOVA followed by Tukey’s HSD).

To establish a genetic relationship between known circadian clock mutants and the long period of *nua-3*, we constructed a series of double and triple mutant lines and assessed their circadian period. In all instances but one, *nua-3* in combination with *cca1lhy* or with the *prr3, prr5, prr7, prr9* or the *prr7 prr9* double mutants exhibited additivity in circadian period (Fig. 3A,B). Strikingly, the short period *toc1-101* mutant was fully epistatic to *nua-3*, with the *nua-3 toc1-101* period identical to *toc1-101* (Fig. 3C).

**Figure 3.**
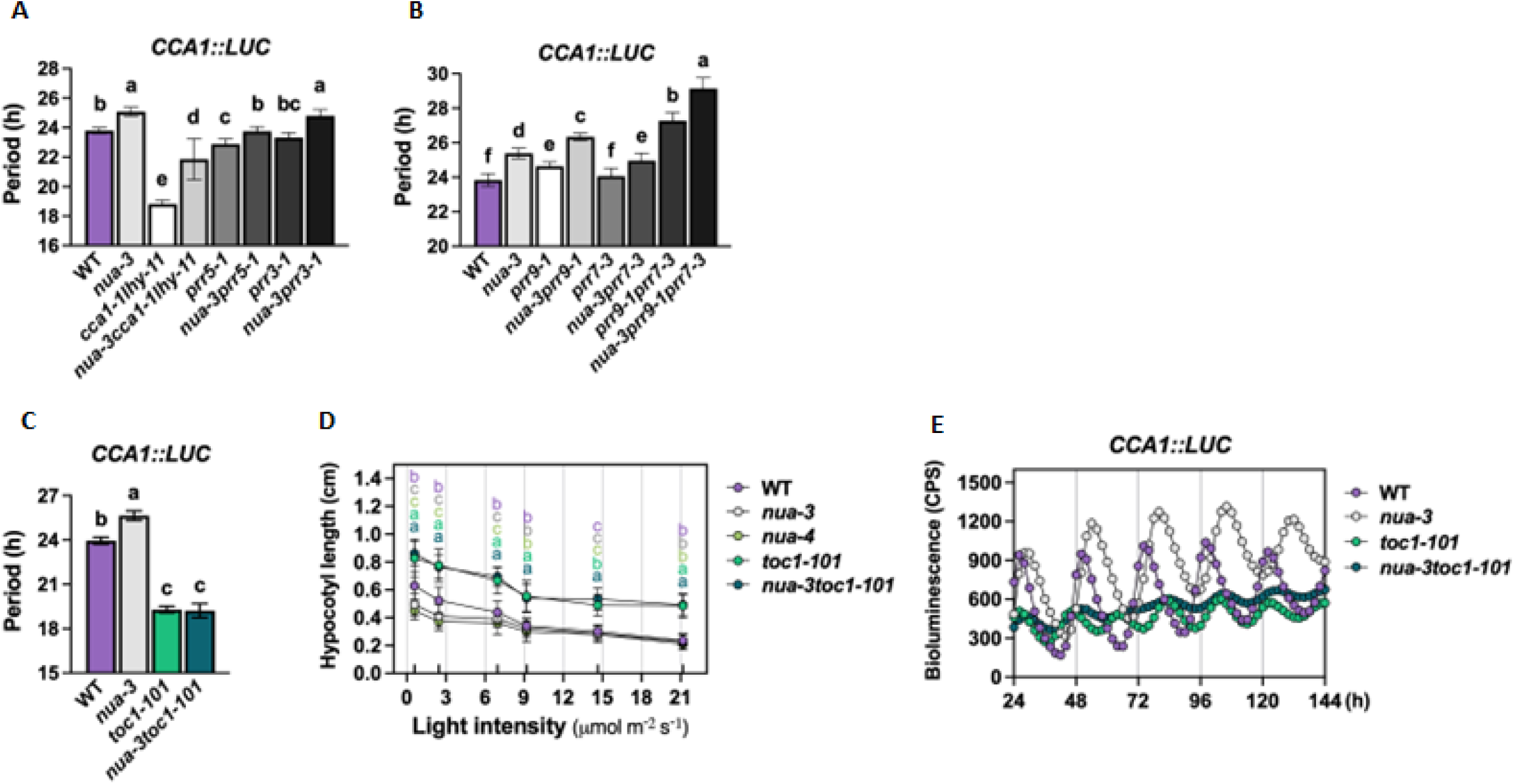
*TOC1* is epistatic to *NUA* in the regulation of circadian period, oscillation amplitude and hypocotyl elongation. **(A, B)** Circadian period estimates of *CCA1:LUC* bioluminescence rhythms under continuous light in *nua-3* single and double mutant combinations with (A) *cca1-1 lhy-11*, *prr5-1*, and *prr3-1*, and (B) *prr9-1*, *prr7-3*, and *prr9-1 prr7-3*. Data are presented as mean ± SD (n ≥ 20). A representative result from two or three biological trials is shown. **(C)** Circadian period estimates of *CCA1:LUC* bioluminescence rhythms under continuous light in WT, *nua-3*, *toc1-101*, and *nua-3 toc1-101*. Data are presented as mean ± SD (n ≥ 23). A representative result from three biological trials is shown. **(D)** Hypocotyl length measurements of WT, *nua-3*, *nua-4*, *toc1-101*, and *nua-3 toc1-101* seedlings grown under continuous light at the indicated intensities. Data are presented as mean ± SD (n ≥ 34). Statistical significance was determined by one-way ANOVA with Tukey’s post hoc test. A representative result from three biological trials is shown. **(E)** *CCA1:LUC* bioluminescence traces under continuous light in WT, *nua-3*, *nua-4*, *toc1-101*, and *nua-3 toc1-101*. Data are presented as mean ± SEM (n ≥ 23). A representative result from three biological trials is shown. Statistical significance was determined by one-way ANOVA with Tukey’s post hoc test.

We further tested genetic epistasis for hypocotyl length. The long hypocotyl of *toc1* mutants ^37^ contrasts with the short hypocotyl *nua* mutant (Fig. 3D). *nua-3 toc1-101* plants are similarly long hypocotyl (Fig. 3D), indicating the need for TOC1 to obtain the short hypocotyl *nua-3* phenotype. Additionally, the low amplitude *CCA1:LUC* oscillations in *nua-3 toc1-1* mutants are very similar to *toc1-101*, in contrast to the very high amplitude *nua-3* oscillations (Fig. 3E). Taken together, these results demonstrate that TOC1 is required for all three *nua* phenotypes (i.e. circadian period, hypocotyl length and *CCA1:LUC* amplitude).

Long period and short hypocotyls are consistent with high levels of TOC1 ^37^. We next examined TOC1-YFP (TMG ^38^) levels in *nua-3* over a 12 hr light/12 hr dark (LD) cycle and found no detectable differences from WT (Fig. 4A,B). As TOC1 is a transcriptional regulator, we considered that TOC1 nuclear levels are likely the important feature in its function. We examined the TOC1 nuclear/cytoplasmic ratio in *nua-3* and *nua-4* mutants and found a disproportionately higher nuclear level of TOC-YFP at ZT 13 and ZT17, time points when TOC1 levels are at their highest (Fig. 4C; Supplementary Fig. 1). Additionally, we quantitated the nuclear/cytoplasmic ratio of TOC1-YFP in living WT and *nua-3* roots by fluorescence imaging and confirmed a higher nuclear/cytoplasmic ratio of fluorescence levels in the *nua-3* background (Fig. 4D). To test for specificity, we examined the effect of the *nua* mutation on the PRR3-GFP nucleocytoplasmic ratio and found no difference from WT (Supplementary Fig. 2). This indicates that the *nua* mutation does not generically alter protein distribution differences between the nucleus and cytosol.

**Figure 4.**
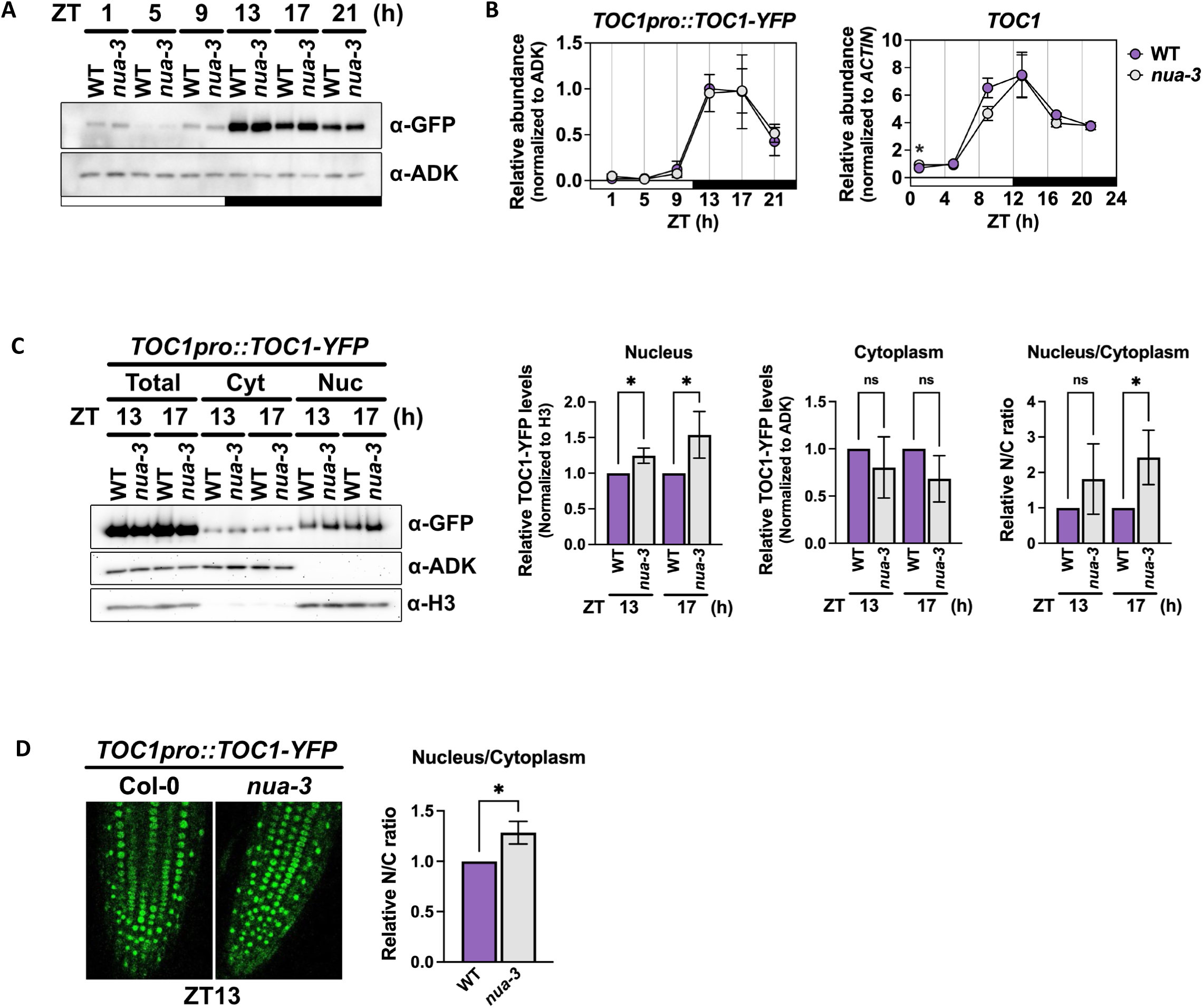
NUA alters TOC1 nucleocytoplasmic distribution. **(A)** Immunoblot analysis of total TOC1 protein levels in WT and *nua-3* plants over a 12/12 L/D time course. TOC1 protein abundance was quantified using *TOC1pro::TOC1-YFP* and normalized to ADK. A representative result from three biological trials is shown. **(B)** Quantification of TOC1 protein levels by immunoblot in WT and *nua-3* plants under 12/12 L/D conditions. Left, TOC1-YFP abundance was normalized to ADK. Right, *TOC1* transcript normalized to *ACTIN*. Data are presented as mean ± SD (n = 2 or 3). Statistical significance was determined by Student’s t-test. **(C)** Subcellular fractionation of TOC1 protein from WT and *nua-3* plants under 12/12 L/D conditions. Bar graphs show the relative TOC1 abundance in the nucleus, cytoplasm, and the nucleus-to-cytoplasm ratio. Data are presented as mean ± SD (n = 3). Statistical significance was determined by Student’s t-test. A representative result from three biological trials is shown. **(D)** Quantification of the nucleus-to-cytoplasm ratio of TOC1 protein in WT and *nua-3* plants by confocal microscopy under 12/12 L/D conditions. Data are presented as mean ± SD (n = 3). Statistical significance was determined by Student’s t-test. A representative result from three biological trials is shown.

### Proteasome-dependent nuclear TOC1 degradation is reduced in *nua*

One plausible reason for the distribution difference is an increased nuclear stability of TOC1. We tested this possibility by inhibiting protein translation using cycloheximide (CHX)-treated seedlings harvested at 3-hour intervals over 12 hours and probing for TOC1-GFP levels in nuclear and cytosolic fractions in WT and *nua-3* plants. Relative to the turnover rate in WT, only the nuclear fraction of TOC1-YFP showed a significantly longer half-life in the *nua* background, indicating an increase in the stability of nuclear TOC1 in the *nua* background (Fig. 5A). When treated with a proteasome inhibitor (MG132) or MG132+CHX, the TOC1-YFP levels were similar in both backgrounds and in both compartments, indicating that nucleocytoplasmic protein translocation remains unchanged by the *nua* mutation and that proteasome-dependent degradation is slower in *nua-3* (Fig. 5B,C).

**Figure 5.**
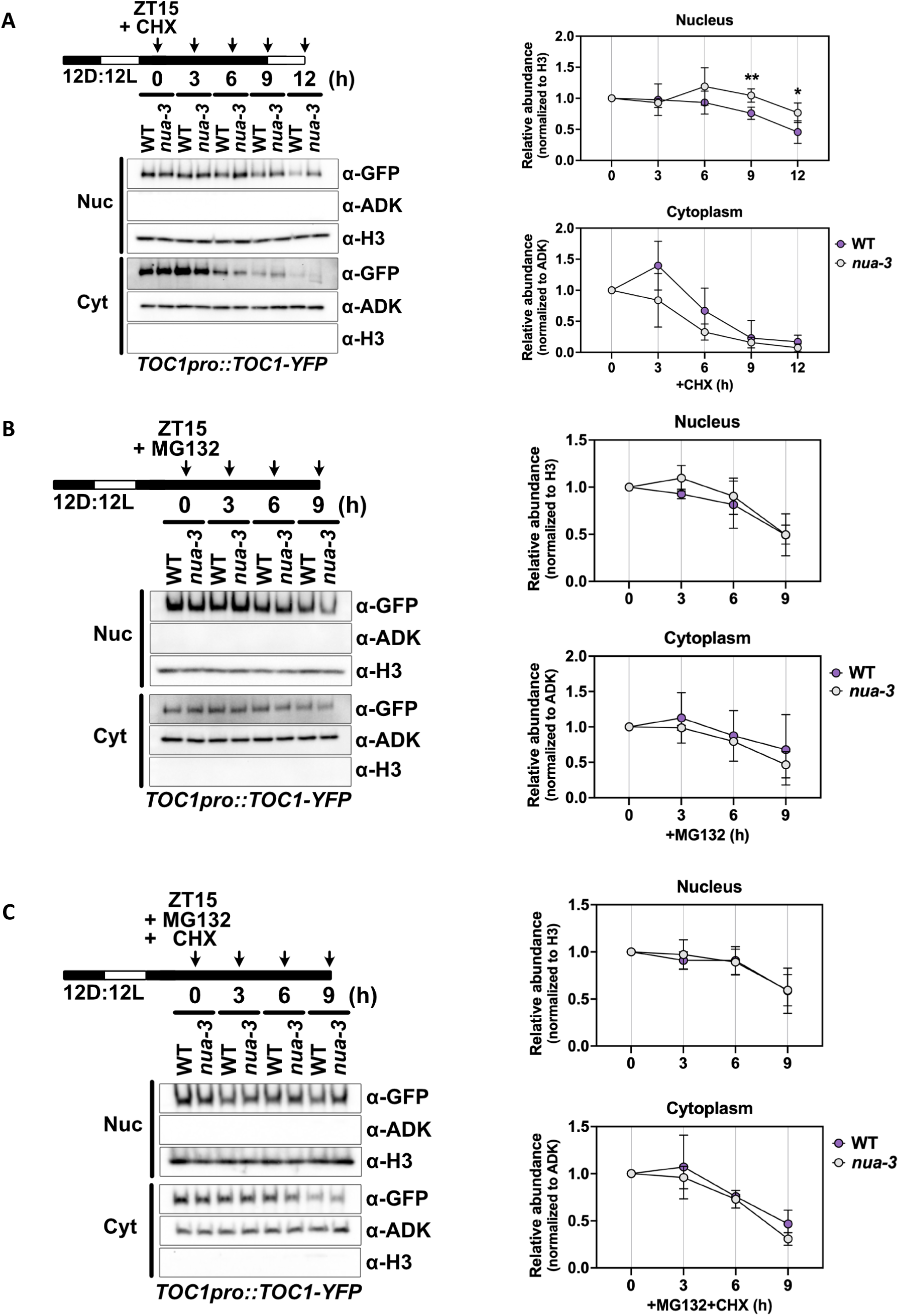
Loss of *NUA* reduces TOC1 protein turnover in the nucleus. (A) TOC1 protein stability in nuclear and cytoplasmic fractions of WT and *nua-3* plants following cycloheximide (CHX) treatment at ZT15. Left, representative immunoblots probed with anti-GFP (TOC1pro::TOC1-YFP), anti-ADK, and anti-H3. Right, quantification of TOC1 abundance in the nucleus and cytoplasm normalized to H3 and ADK, respectively. A representative result from three biological trials is shown. (B) TOC1 protein levels in nuclear and cytoplasmic fractions of WT and *nua-3* plants following MG132 treatment. Immunoblotting and quantification were performed as in (A). A representative result from three biological trials is shown. (C) TOC1 protein levels in nuclear and cytoplasmic fractions of WT and *nua-3* plants following combined MG132 and CHX treatment. Immunoblotting and quantification were performed as in (A). A representative result from three biological trials is shown.

### The *nup136* mutant phenocopies *nua*

These findings suggested aberrant proteasome function, specific to the nucleus, in the *nua* background. Previous IP-MS work has linked the nuclear basket protein, NUP136 (also referred to as NUP1^35^), with various proteasome subunits and NUA ^39^. Additionally, a recent report showed that the NUP136 ortholog in mouse, NUP153, anchors the NUA ortholog, Tpr, to the murine nuclear basket ^40^. With this in mind, we first tested NUP136-NUA interactions through yeast two hybrid and co-IP assays, using NUA polypeptide fragments and designations previously described ^31^. N-terminal (F1R2) and C-terminal (CT) regions of NUA were tested separately due to the large size of the full-length protein. Both assays showed N-terminal NUA interacts more strongly with NUP136 than the C-terminal region (Fig. 6A, B). Consistent with the notion of the anchoring and stabilization of NUA to the nuclear basket through association with NUP136, NUA protein levels are significantly diminished in the *nup136-1* mutant (Fig. 6C). In contrast, NUP136-GFP levels are unaffected in the *nua-3* background (Fig. 6D).

**Figure 6.**
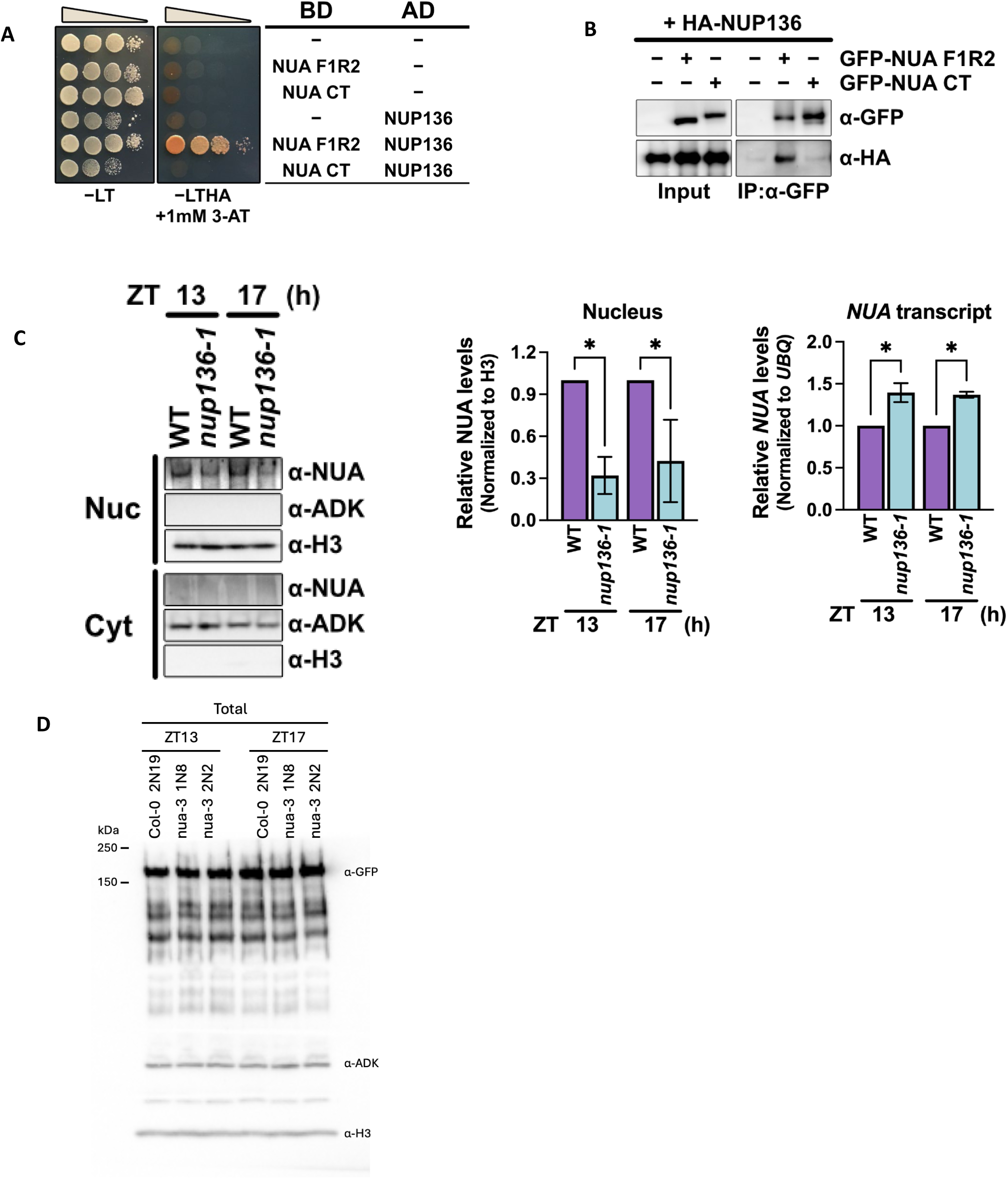
NUA is stabilized through interaction with NUP136. **(A)** Yeast two-hybrid analysis of the interaction between NUP136 and NUA. Yeast cells co-transformed with the indicated bait and prey constructs were plated on selective medium (–Leu–Trp–His–Ade + 1 mM 3-AT) and grown for 4 days. **(B)** *In vivo* co-immunoprecipitation of NUP136 and NUA. HA-NUP136 and GFP-NUA were transiently co-expressed in leaves of 3-week-old *Nicotiana benthamiana*, and total protein extracts were subjected to immunoprecipitation using an anti-GFP antibody. The resulting immunocomplexes were analyzed by immunoblotting with anti-GFP and anti-HA antibodies. Five independent experiments were conducted. **(C)** Left, immunoblot analysis of NUA protein levels in nuclear and cytoplasmic fractions of WT and *nup136-1* plants at ZT 13 and ZT 17. Immunoblots were probed with anti-NUA, anti-H3 (nuclear marker), and anti-ADK (cytoplasmic marker). Middle, quantification of NUA protein levels in the nucleus normalized to H3. Right, RT-qPCR analysis of *NUA* transcript levels normalized to *UBQ*. Data are presented as mean ± SD (n = 3). A representative result from three biological trials is shown. **(D)** Immunoblot analysis of NUP136-GFP protein levels in nuclear and cytoplasmic fractions of WT and *nua* plants. Immunoblots were probed with anti-GFP (*NUP136pro::NUP136-GFP*), anti-H3 (nuclear marker), and anti-ADK (cytoplasmic marker).

Further experiments with transgenic *NUP136pro::NUP136-GFP* expressing Arabidopsis indicates a functional relationship between NUP136 and NUA *in vivo*. NUP136-GFP localizes evenly along the nuclear rim, as expected for a nuclear basket protein (Fig. 7A). In *nua-3* the smoothly even NUP136-GFP signal is disrupted and distinct puncta are seen both in single-z sections and in 3D reconstruction (Fig. 7B, C; Supplementary Fig. 3). In contrast, the nuclear rim localization patterns of the outer ring nucleoporin RAE1 and the nuclear membrane protein WIP1 are not affected by the absence of NUA (Fig. 7D). These results suggest that a specific NUP136/NUA stoichiometry is required for appropriate positioning of NUP136 to the nuclear pore. At this level of resolution we cannot determine if the overall nuclear pore density or structure has been altered by *nua-3*.

**Figure 7.**
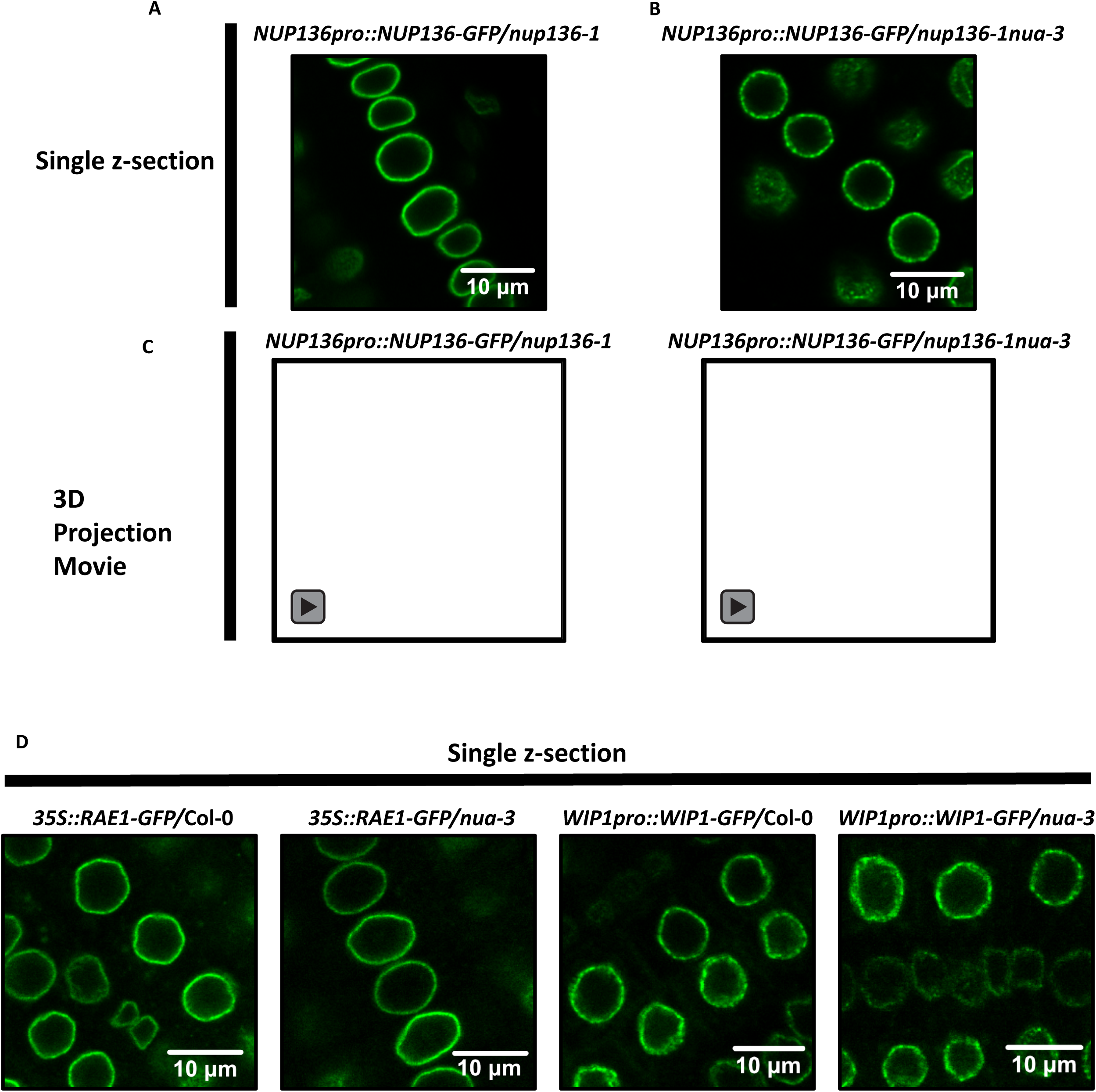
NUP136-GFP nuclear envelope localization is altered in *nua* mutants. Confocal microscopy images of *NUP136pro::NUP136-GFP* in 5-day-old roots of (A) *nup136-1* and (B) *nup136-1 nua-3* in single z-section images. (C) 3D-reconstructed images generated from serial confocal z-sections. Scale bars are indicated. (D) Confocal microscopy single z-section images of 5-day-old roots expressing *35S::RAE1-GFP* (left) or *WIP1pro::WIP1-GFP* (right) in Col-0 and *nua-3* backgrounds. Scale bars are indicated.

We considered that the NUA-NUP136 interactions demonstrated in these assays could reflect an *in vivo* association at the nuclear basket. If so, *nup136* mutants might exhibit similar circadian and hypocotyl phenotypes as *nua* mutants. Testing this, we found that like *nua-3*, *nup136-1* is long period (*nup136-1*: 25. 7; WT: 24.2) and the *toc1-101 nup136-1* double mutant period (20.5 hrs) is very similar to *toc1-101* (19.7 hrs)(Fig. 8A). Additionally, the long hypocotyl of *toc1-101* is epistatic to the shorter hypocotyl of *nup136-1* in the *toc1-101 nup136-1* double mutant (Fig. 8B), similar to the relationship seen between *nua-3* and *toc1-101*(Fig. 3B). These findings demonstrate that *nua* and *nup136* mutants similarly require TOC1 for their period and hypocotyl phenotypes, and together with their physical interaction, they likely form a functional complex at the nuclear basket *in vivo*.

**Figure 8.**
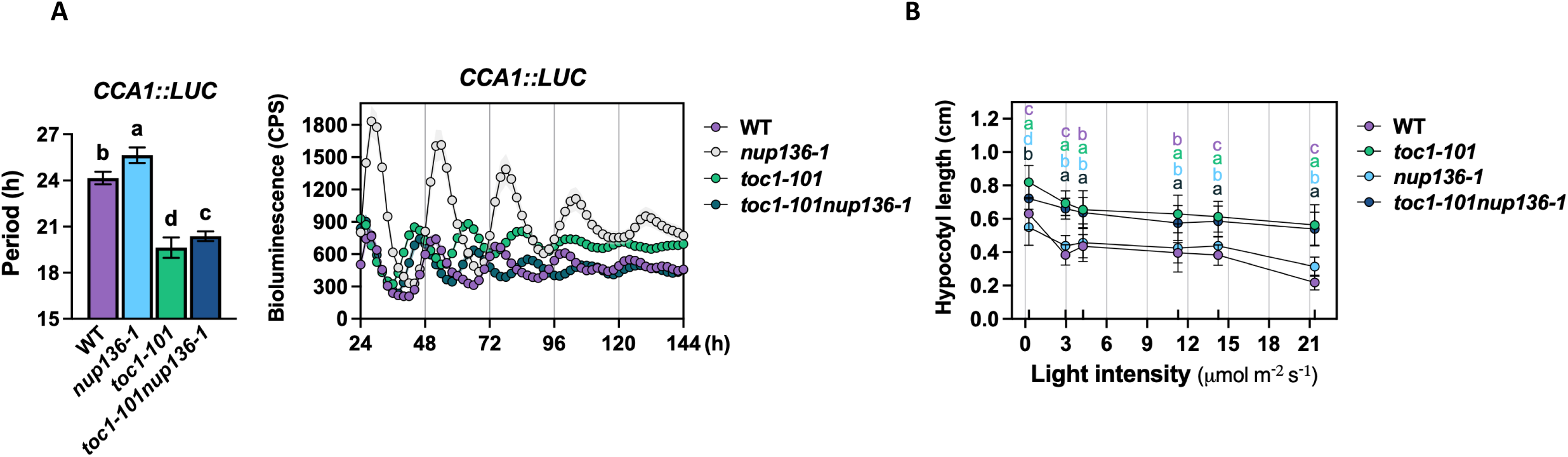
Genetic interaction between *NUP136* and *TOC1* in the regulation of circadian clock function. **(A)** Circadian period length analysis in WT, *nup136-1*, *toc1-101*, and *toc1-101 nup136-1* plants under constant light conditions. Data are presented as mean ± SD. Statistical significance was determined by one-way ANOVA with Tukey’s post hoc test. **(B)** Measurement of hypocotyl length in WT, *nup136-1*, *toc1-101*, and *toc1-101 nup136-1* seedlings grown under continuous light for 4 days. Data represent mean ± SEM (n = 12–24 seedlings per genotype). A representative result from three independent experiments is shown.

### Proteasome-dependent degradation of TOC1 depends on NUP136 and NUA N-terminus dosage

In support of this notion, we next tested TOC1-YFP levels, after CHX treatment, in the nuclear and cytoplasmic fractions in the *nup136-1* background, as done with *nua-3 (*Fig. 5A). In *nup136-1* TOC1-YFP levels were more stable in the nuclear fraction only, indicating that NUP136 is required for normal nuclear TOC1 turnover (Fig. 9A), as found with *nua-3*. When crossed into the *nup136-1* background *TMG* expression was reduced significantly (Supplementary Fig. 4), likely from some degree of *TMG* transgene silencing, which accounts for the lower protein levels relative to WT (Fig. 9A). However, this does not affect the protein turnover rates as measured by the CHX chase assay.

**Figure 9.**
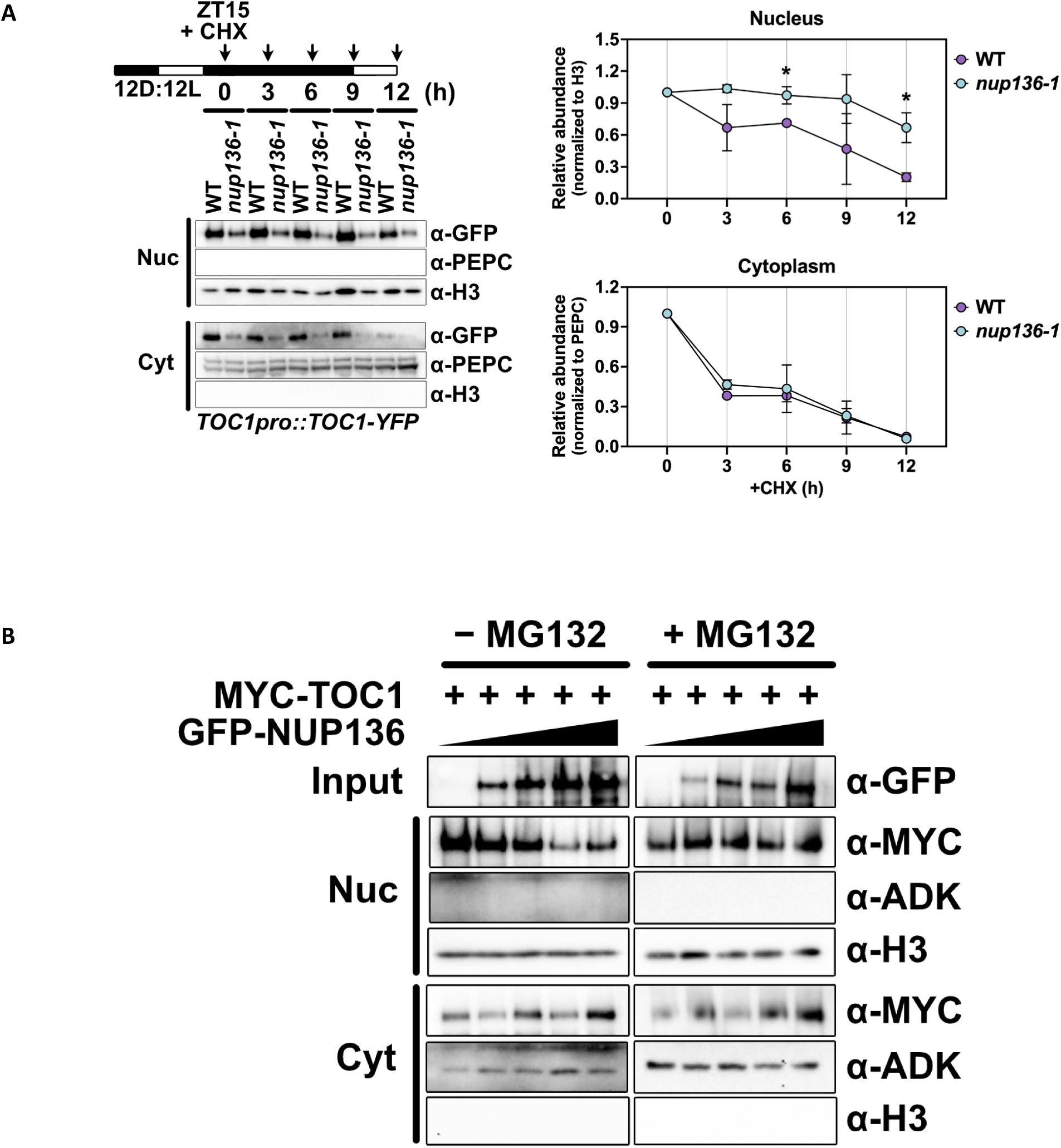
NUP136 dosage determines TOC1 nuclear abundance. **(A)** Subcellular fractionation of TOC1-YFP protein in WT and *nup136-1* plants. Left, a representative immunoblot of nuclear and cytoplasmic fractions probed with anti-GFP (TOC1pro::TOC1-YFP) from three biological trials. Right, quantification of TOC1-YFP protein levels in the nucleus and cytoplasm, normalized to H3 and PEPC, respectively. Data are presented as mean ± SD (n = 3). **(B)** Subcellular fractionation of TAP-TOC1 protein in nuclear and cytoplasmic fractions upon increasing expression levels of GFP-NUP136 in the absence (left) or presence (right) of MG132 treatment. Immunoblots were probed with anti-GFP (GFP-NUP136), anti-MYC (TAP-TOC1), anti-ADK (cytoplasmic marker), and anti-H3 (nuclear marker). A representative result from three biological trials is shown.

Conversely, we transiently overexpressed NUP136 together with myc-TOC1 in *N. benthamiana* to assess the effects of excess NUP136 on TOC1 levels in the cytosol and nucleus. Using different infiltration titers, we adjusted the relative levels of the two proteins and found that increasing expression of NUP136 diminished the nuclear abundance of myc-TOC1 but had no effect on the cytosolic levels (Fig. 9B). When co-infiltrated with MG132 nuclear myc-TOC1 levels were unaffected by higher NUP136 expression (Fig. 9B), indicating proteasome-dependent turnover is responsible for lower myc-TOC1 accumulation.

We performed similar experiments using the N-terminus and C-terminus of NUA to test effects on TOC1 levels. As the NUA nuclear localization signal (NLS) is at the C-terminus^31^ we included an NLS to F1R2 to ensure nuclear presence. When we expressed increasing levels of a GFP-NLS-NUA F1R2 protein with myc-TOC1 only the nuclear fraction of TOC1 was concomitantly reduced (Fig. 10A), consistent with the nuclear localization of the N-terminal portion of NUA (Fig. 10B). In contrast, myc-TOC1 levels were unaffected by increasing levels of nuclear C-terminal NUA (Fig. 10 C, D).

**Figure 10.**
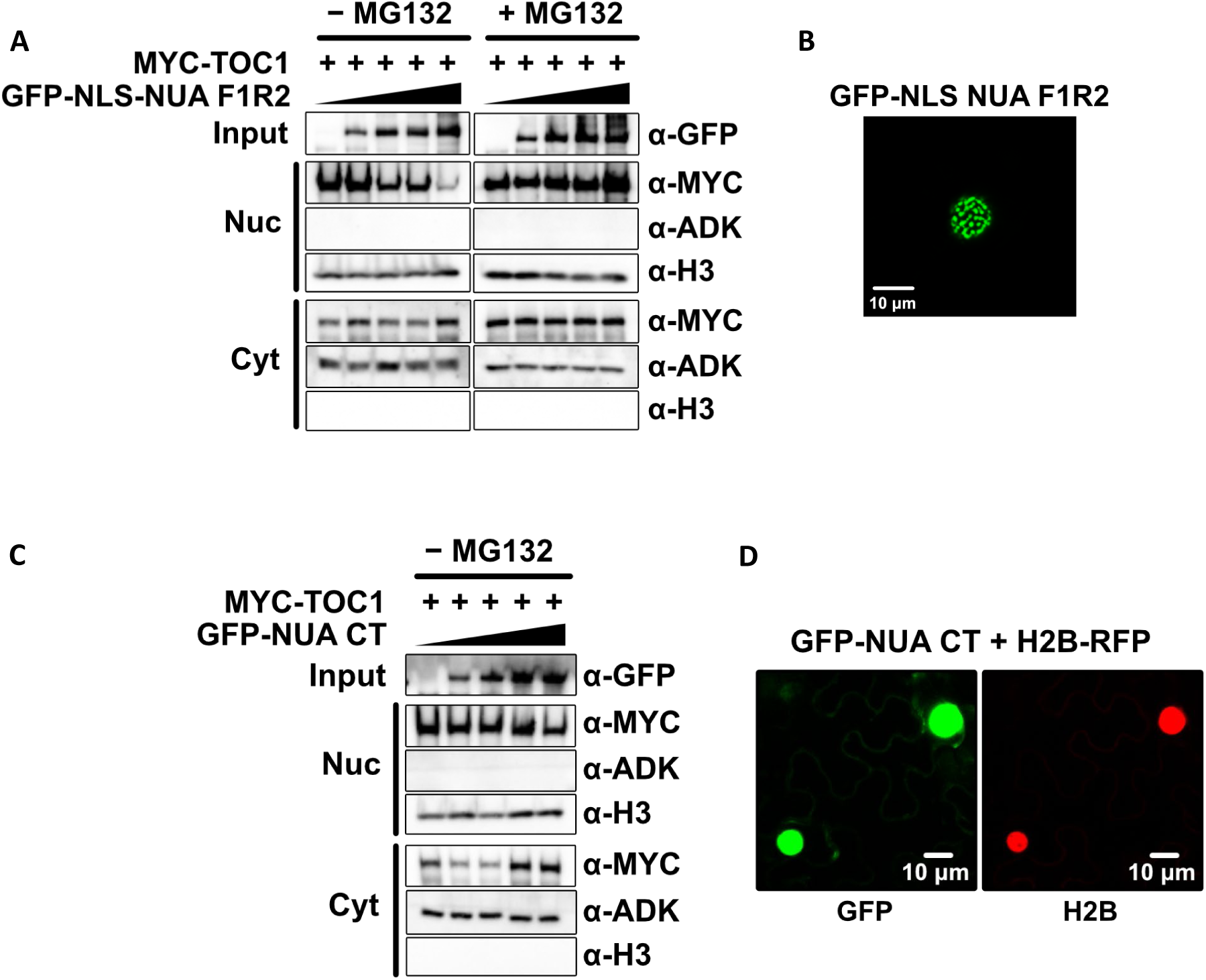
NUA N-terminus promotes nuclear depletion of TOC1 (. **(A)** Left, subcellular fractionation and immunoblot analysis of TAP-TOC1 protein in nuclear and cytoplasmic fractions upon increasing expression levels of GFP-NLS-NUA F1R2 in the absence or presence of MG132 treatment. Immunoblots were probed as in (A). A representative result from three biological trials is shown. Right, nuclear localization of GFP-NLS-NUA F1R2 under confocal microscopy. **(B)** Confocal microscopy images of GFP-NLS-NUA F1R2. Scale bars are indicated. **(C)** Subcellular fractionation and immunoblot analysis of TAP-TOC1 protein in nuclear and cytoplasmic fractions upon increasing expression levels of GFP-NUA CT. Immunoblots were probed as in (A). A representative result from three biological trials is shown. **(D)** Subcellular localization of GFP-NUA CT in *N. benthamiana* leaves co-infiltrated with H2B-RFP as a nuclear marker, visualized by confocal microscopy.

These results show that TOC1 abundance is inversely correlated with levels of NUP136 and the NUA N-terminus. Taken together with the CHX experiments these findings strongly implicate an association of these two nuclear basket proteins with nuclear proteasome activity.

### NUP136 re-localizes the NUA N-terminus and proteasome components to the inner nuclear periphery *in vivo*

To further understand the relationships between NUP136, NUA and TOC1 at the nuclear basket we tested interactions between TAP-TOC1 and HA-NUP136 and the NUA N-and C-termini tagged with GFP. Transient co-expression in *N. benthamiana* of these proteins alone and in pairwise and triple combinations, followed by immunoprecipitation of TOC1 established that only NUP136 co-immunoprecipitates (Fig. 11A). However, when retested with a nuclear-localized NUA N-terminus (GFP-NLS-NUA F1R2) TOC1 was able to associate with the NUA N-terminus (Fig. 11B). This suggests that NUA nuclear localization is required for a TOC1-NUA interaction, possibly drawing on the presence of the endogenous NUP136 present in the nuclear basket of *N. benthamiana*.

**Figure 11.**
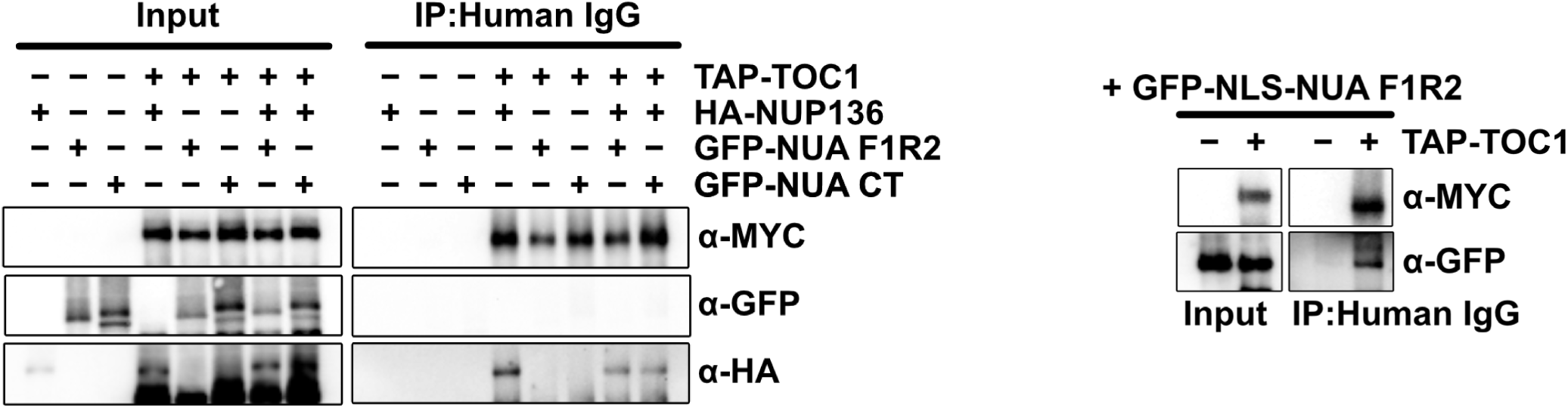
Co-immunoprecipitation analysis of the interaction between NUP136, NUA, and TOC1. **(A)** Co-immunoprecipitation (Co-IP) assay examining the interaction among TAP-TOC1, HA-NUP136, GFP-NUA F1R2, GFP-NUA CT, and GFP-NLS-NUA F1R2 in the indicated combinations. The proteins were transiently co-expressed in *N. benthamiana* leaves via *Agrobacterium*-mediated infiltration. Protein extracts were immunoprecipitated with human IgG and immunoblotted with anti-MYC (TAP-TOC1), anti-HA (HA-NUP136), and anti-GFP (GFP-NUA F1R2, GFP-NUA CT, or GFP-NLS-NUA F1R2). Input samples are shown on the left. A representative result from three biological trials is shown. **(B)** Co-IP assay testing the interaction between TAP-TOC1 and GFP-NLS-NUA F1R2. Protein extracts were immunoprecipitated with anti-GFP antibody and immunoblotted with anti-MYC and anti-GFP

We next examined the *in vivo* relationships between NUA, NUP136, TOC1 and the proteasome using fluorescent confocal microscopy to assess nuclear and subnuclear localization of each component, and their effects on nuclear positioning of each other. Transient expression of NUP136-CFP in *N. benthamiana* cells regularly showed fluorescence along the inner nuclear periphery, recapitulating the pattern observed in stably transformed NUP136::NUP136-GFP (compare Fig. 12 and Fig. 7A; Supplementary Fig. 5: 3D reconstruction). In contrast NUA-F1R2-NLS-YFP formed fluorescent intranuclear bodies that varied with each cell but were typically absent from the nuclear rim (Fig. 12A). NUA CT-YFP filled the nucleus with a diffuse fluorescent signal, exclusive of the nucleolus (Fig. 12A). We additionally tested two components of the proteasome, RPT5b-YFP and RPN1a-YFP. Fluorescent signals for both proteins were present in the cytosol and diffuse within the nucleus, also excluded from the nucleolus (Fig. 12A).

**Figure 12.**
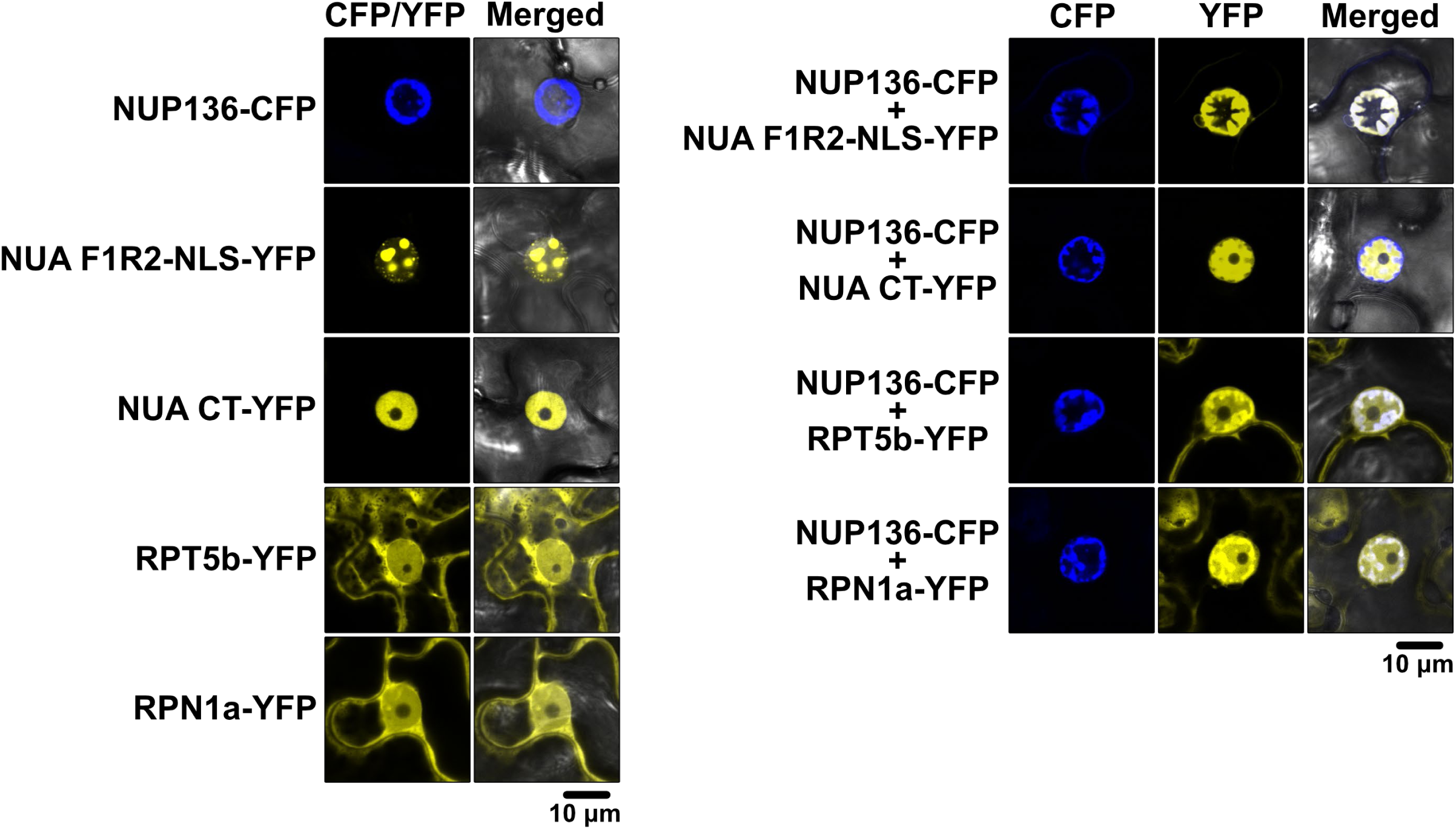
Colocalization analysis of NUP136 with NUA fragments and proteasome subunits in *N. benthamiana* leaves. Confocal microscopy images of *N. benthamiana* leaf epidermal cells transiently expressing the indicated constructs at 3 days post-infiltration. Left, single expression of NUP136-CFP, NUA F1R2-NLS-YFP, NUA CT-YFP, RPT5B-YFP, and RPN1A-YFP. Right, co-expression of NUP136-CFP with NUA F1R2-NLS-YFP, NUA CT-YFP, RPT5B-YFP, or RPN1A-YFP. CFP, YFP, and merged channels are shown. Scale bars are indicated.

We next tested pairwise co-expression of NUP136-CFP with each of these 4 YFP-tagged proteins. The N-terminus of NUA (NUA-F1R2-NLS-YFP) regularly re-localized to the nuclear rim when co-expressed with NUP136-CFP, coincident with position of NUP136-CFP (Fig. 12B). This finding is consistent with yeast 2-hybrid and co-IP assays showing interaction (Fig. 6) and supports the notion that NUP136 can recruit the N-terminus of NUA to its subnuclear position. When HA-NUP136 was co-expressed with NUA-F1R2-CFP, NUA-F1R2 was similarly re-localized to the nuclear rim (Supplementary Fig. 6). These results are consistent with NUP136 recruitment of N-terminal NUA and demonstrate that the nature of the protein tag is not a factor in NUP136-NUA interactions and NUA subnuclear repositioning.

In contrast, when co-expressed with NUP136-CFP the position of NUA CT-YFP remained diffusely present throughout the nucleoplasm and possibly excluded from the nuclear rim position of the co-expressed NUP136-CFP (Fig. 12B; Supplementary Fig. 7)). This result aligns with the absence of a NUP136-NUA C-terminus interaction seen *in vitro* and two-hybrid tests (Fig. 6) and indicates that NUP136 and NUA interact via the N-terminal portion of NUA.

Interestingly, the patterns of both proteasome components (RPT5b-YFP, RPN1a-YFP) showed enhanced yellow fluorescence coincident with the expression position of NUP136-CFP (Fig. 12B; Fig. 8). Similar re-localization of both proteasome components was observed when co-expressed with HA-NUP136, but not when co-expressed with HA-NUA CT (Supplementary Fig. 9A,B,C). These results are consistent with the recruitment of a portion of the nuclear proteasome to the subnuclear nuclear position of NUP136. As a negative control we co-expressed RH6-CFP, a DEAD Box helicase^41^, with both RPN1a-YFP and RPT5b-YFP and found no effect of RH6 expression on the nuclear distribution of these proteasome components (Supplementary Fig. 10).

We repeated these experiments with the NUA N-terminus (NUA F1R2-NLS-YFP). Neither RPN1a nor RPT5b subnuclear localization was altered when co-expressed (Fig. 13; Supplementary Fig. 11), indicating NUA is not involved in proteasome positioning to the nuclear basket.

**Figure 13.**
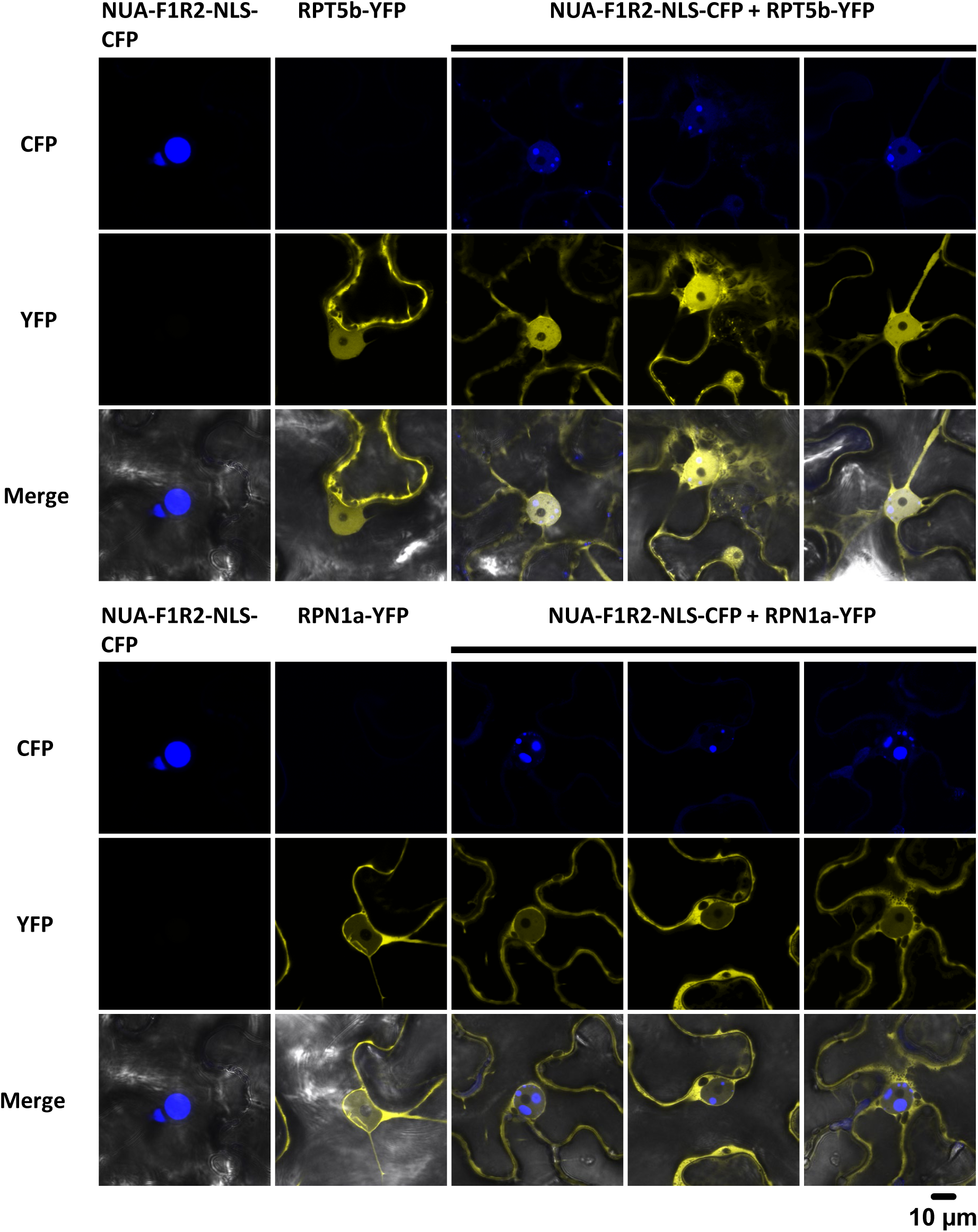
NUA N-terminus does not colocalize with RPT5B-YFP and RPN1A-YFP in *N. benthamiana* leaves. Confocal microscopy images of *N. benthamiana* leaf epidermal cells transiently expressing the indicated constructs at 3 days post-infiltration. Above, co-expression of RPN1a-YFP and NUA-F1R2-NLS-CFP. Below, co-expression of RPT5B-YFP with NUA-F1R2-NLS-CFP. CFP, YFP, and merged channels are shown. Scale bars are indicated.

### NUP136 and NUA N-terminus can re-localize TOC1 to their subnuclear positions

Using these same co-localization assays, we next asked whether these two nuclear basket proteins can associate with TOC1 *in vivo*. Transient expression of TOC1-YFP generally exhibited an apparently random distribution of small nuclear speckles, though in some cases larger yellow inclusions were observed (Fig. 14A). Coexpression with NUP136-CFP or HA-NUP136 relocalized TOC1-YFP to the inner nuclear periphery (Fig. 14B,C; Supplementary Fig. 12). These *in vivo* results support the co-IP assays indicating a TOC1-NUP136 interaction (Fig. 10). Additionally, they demonstrate a novel role for NUP136 in recruiting TOC1 to the nuclear periphery. Similarly, the NUA N-terminus (NUA F1R2-NLS-CFP) co-localized with TOC1-YFP (Fig.14D), but since both proteins are nucleoplasmic no distinctive change in either protein’s position is apparent. Similar experiments with co-expression of NUP136 and PRR5-YFP and ELF4-GFP did not re-localize those proteins (Supplementary Fig. 13).

**Fig 14.**
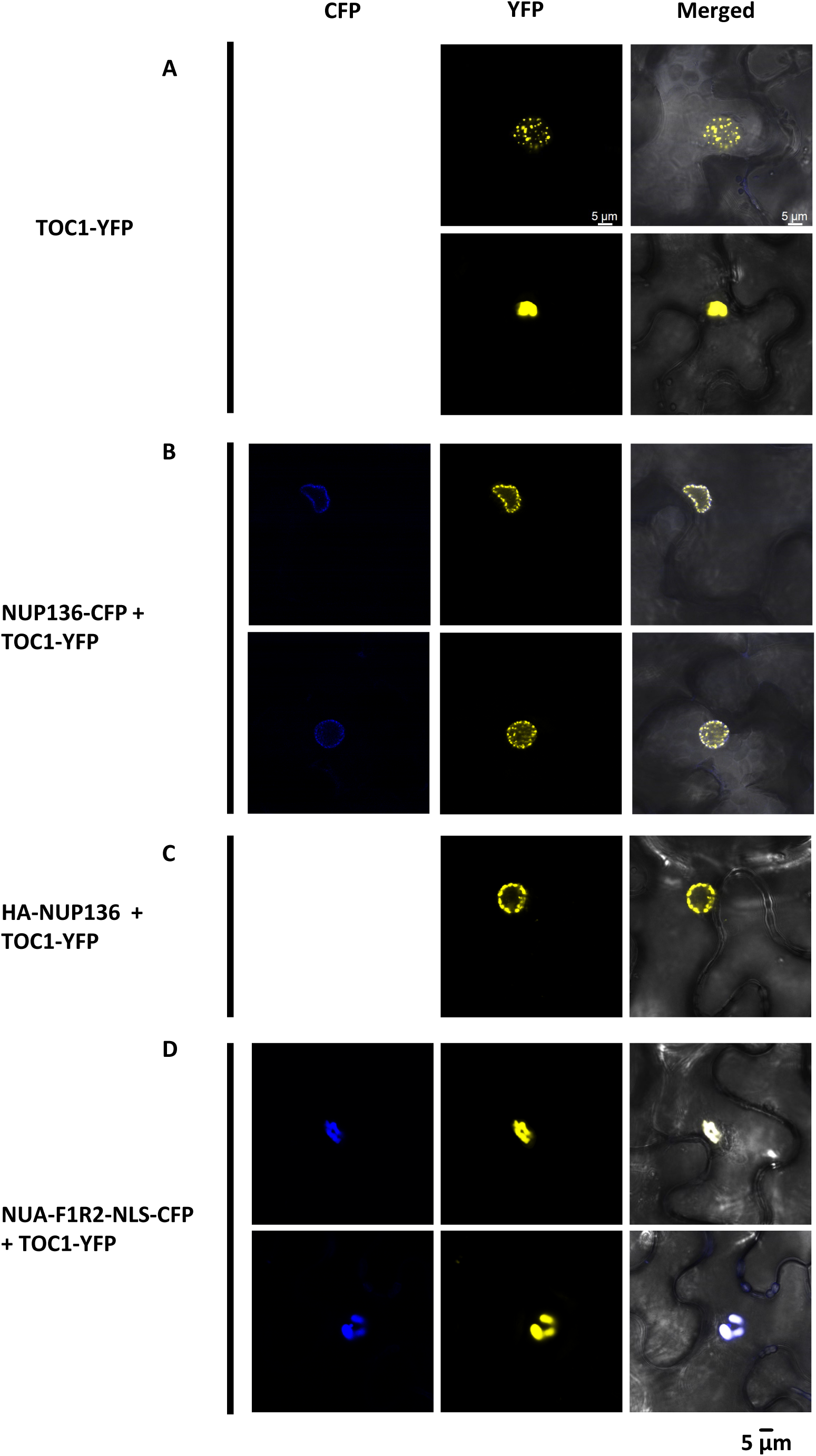
NUP136 and NUA-NT can relocalize TOC1 subnuclear position. Confocal microscopy images of N. benthamiana leaf epidermal cells transiently expressing the indicated constructs at 3 days post-infiltration. **(A)** Two patterns of TOC1-YFP subnuclear expression. **(B)** TOC1-YFP co-expressed with NUP136-CFP or **(C)** HA-NUP136. **(D)** TOC1-YFP co-expressed with NUA-F1R2-NLS-CFP. Scale bars are indicated.

Taken together, the co-localization results support a model in which a NUP136-NUA nuclear basket complex anchors a portion of the nuclear-localized proteasome which facilitates nuclear TOC1 degradation when in association with these basket components (Fig. 15). The long periods of the *nua* and *nup136* mutants result from aberrantly higher TOC1 accumulation through a deficiency in proteasome-dependent TOC1 turnover. Circadian period is highly sensitive to TOC1 dosage (Supplementary Fig. 14), consistent with the notion that slight changes in TOC1 levels are reflected in detectable changes in free-running period.

**Figure 15.**
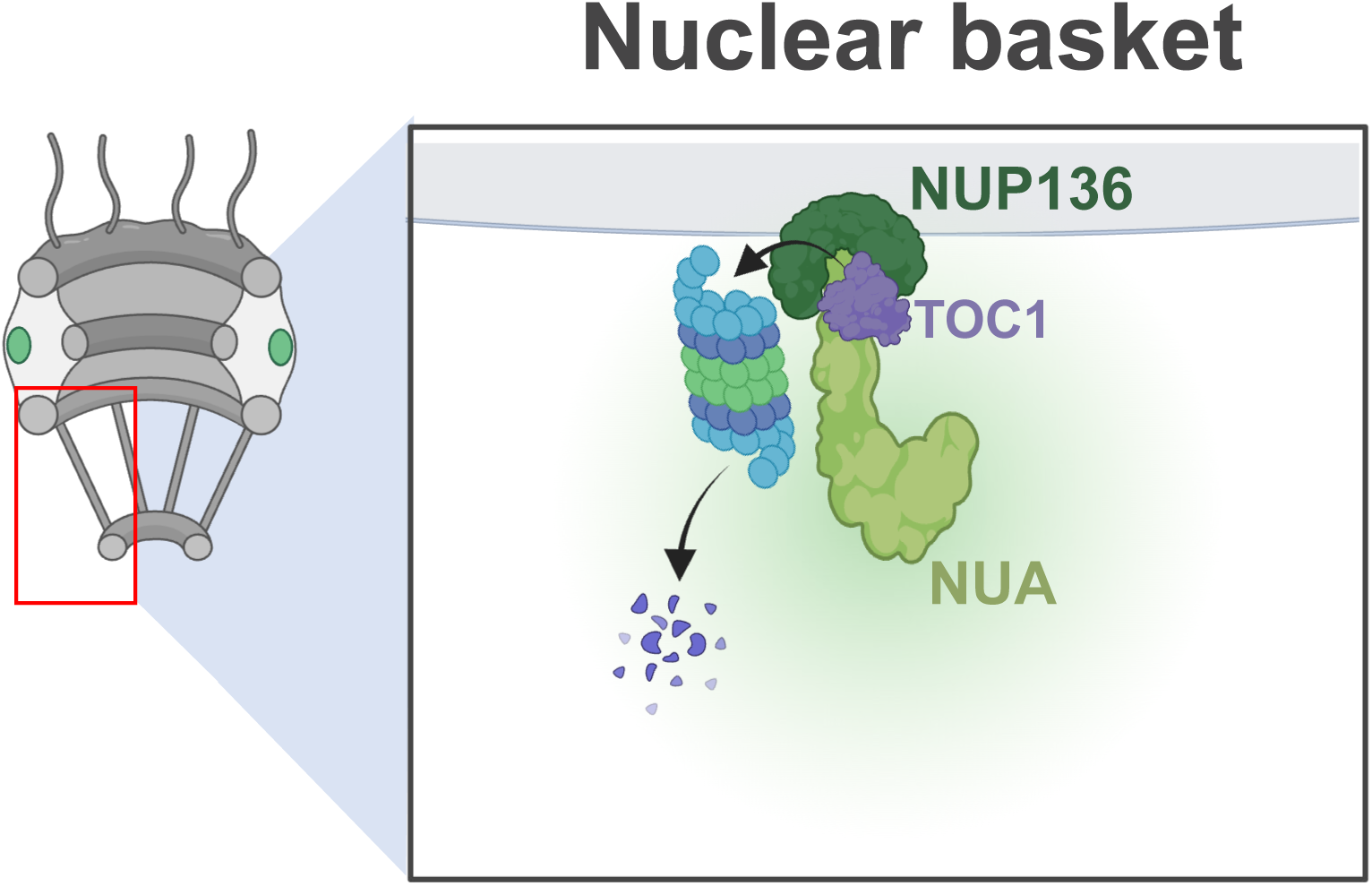
Proposed model for NUP136- and NUA-mediated regulation of TOC1 turnover at the nuclear basket. Schematic illustration of the proposed mechanism by which NUP136 and NUA control TOC1 nuclear abundance. As TOC1 shuttles from the cytoplasm to the nucleus it transits through the nuclear basket. The NUP136-NUA interaction, as part of the nuclear basket complex, creates an environment that facilitates the local residence of 26S proteasomes. TOC1 interaction with the NUP136-NUA complex leads to a proteasome-dependent degradation of TOC1. Loss of *NUP136* or *NUA* disrupts this regulatory environment, leading to aberrant TOC1 accumulation in the nucleus and consequent alterations in circadian period, amplitude, and hypocotyl elongation.

## Discussion

Recent physiological, structural and molecular studies have improved our understanding of the role of the nuclear basket in physiology and development. In animal systems, some human diseases and developmental anomalies have been tied to deficiencies in the function of the nuclear pore, and specifically the nuclear basket ^42–44^. In plants, flowering time, seedling establishment, plant immune responses, auxin responses and the circadian clock are all affected by defects in the nuclear pore and the associated basket ^31,32,45–47^

In this study we have provided a molecular basis for understanding the function of the nuclear basket in circadian regulation by identifying the role of the nuclear basket in the subnuclear positioning of proteasomes. Our findings support a model in which *nua* and *nup136* mutants exhibit long circadian periods due to aberrantly high levels of nuclear TOC1 resulting from reduced nuclear proteolysis. We suggest that a nuclear basket complex, formed in part through a NUP136-NUA interaction, anchors a portion of the nuclear proteasome population to the pore region. Absent either NUP136 or NUA, basket-associated proteasomes are lacking, reducing local proteolytic capability.

This conclusion is based first on our observations of a NUA-NUP136 physical interaction (Fig. 6). Supporting this we found that NUA protein levels are diminished in the *nup136* mutant (Fig. 6), and that transient NUP136 expression can re-localize only the NUA N-terminus to the inner nuclear rim (Fig. 12). This is consistent with recent studies of the nuclear basket in mouse (m) and yeast (y) using cryo-electron tomography to position NUP153(m) and NUP60(y) in association with nuclear pore (Nup) components on the nucleoplasmic side. These proteins then interact with the N-terminus of Tpr(m)/Mlp(y) to form the “struts” of the basket ^40^. Plant NUP136/NUP1 and NUA are the functional homologs of mouse NUP153 and Tpr, respectively^25^.

The importance of a proper balance of NUP136 and NUA in the formation of the nuclear basket is seen in the appearance of discrete NUP136-YFP speckles along the nuclear envelope in the *nua-3* mutant, in contrast to an evenly clear fluorescence in the WT (Fig. 7). *nua-3* is not a null mutation, with some evidence that a small amount of only the N-terminal portion of the protein is made^31^. The *nua-4* null mutant has a more severe developmental and circadian phenotypes (Fig.1; ^31^). Hence, the mislocalization of NUP136 may result from the presence of low levels of the “strut” portion of N-terminus of NUA, and the complete absence of the C-terminal “basket ring” portion of NUA, resulting in an overall disruption in nuclear basket positioning. Additionally, Tpr(m) and Mlp(y) are dimeric in their respective nuclear baskets^40^. Therefore, if NUA is also normally dimeric in the Arabidopsis nuclear basket the absence of the NUA C-terminus in *nua-3* may destabilize the basket and affect NUP136 location. At the same time, NUA protein levels are diminished in the *nup136-1* null mutant (Fig. 6). This result is consistent with the notion that NUP136 is required for anchoring NUA to the nucleoplasmic side of the pore to create the basket and indicates the reciprocal importance of appropriate levels of both proteins for basket location and stability.

We found that only NUP136, not NUA, can interact with and relocalize to the nuclear periphery two components of the regulatory particle (RP) of the 26S proteasome, RPN1a and RPT5b^48^ (Fig. 12-13; Supp. Fig. 9). Both proteins have been previously reported in co-immunoprecipitates with NUP136/NUP1 in Arabidopsis ^39^ and recently other RP components have been nuclear localized with NUP136/NUP1in Arabidopsis^47^. Additionally, work in *S. pombe* showed an absence of proteasome enrichment near the nuclear envelope in the *alm1* mutant, a protein with functional homology to Tpr/MLp1/NUA ^49^. Supporting this localization, cryo-electron tomography in *Chlamydomonas* showed proteasome concentrations at the nuclear pore and envelope^50^. Thus, across yeast, plant and algal systems there is strong evidence for at least a portion of the nuclear proteasome positioned at or near the nuclear pore basket.

TOC1 interacts with both components of the nuclear basket, *in vitro* and *in vivo* (Figs. 11 and 14). Transient TOC1 expression alone is generally distributed in small, dispersed speckles within the nucleus, whereas co-expression with NUP136 relocalizes TOC1 to the nuclear rim. TOC1 co-expression with the NUA N-terminus results in a mutual nucleoplasmic co-localization (Fig. 14). Hence, our results support a physical recruitment of TOC1 to the nuclear basket through an interaction with both core basket elements, NUP136 and NUA.

There is ample evidence for a role for the nuclear pore in the control of gene expression and genome organization in metazoans^30,51,52^. As well, a recent extensive multiomics atlas of the Arabidopsis nuclear envelope has further established networks of interactions that implicated the envelope in an additional wide range of processes, including RNA processing and possibly proteolysis ^29^. More specifically, recent work in Arabidopsis using proximity labelling proteomics has identified roles for GBPL3 (guanylate-binding protein (GBP)-like GTPase), a nuclear basket protein, as capable of recruiting transcriptional regulators and RNA processing proteins ^53^. The absence of proteasome component identification in that work may point to discrete subregions of activity at the basket, with proteasome location and activity sequestered from other basket functions.

However, TOC1, as a transcriptional repressor, may be in close proximity to the basket proteasomes if TOC1-chromatin associations are already present in the vicinity through associations with other basket proteins (e.g. GBPL3). This arrangement could provide for an efficient mechanism for elimination/degradation of transcriptional regulators, contingent on additional conditions.

It remains unclear whether the previously identified specifier of TOC1 degradation, the E3 ligase substrate adaptor F-box protein ZTL, is present in the nucleus in sufficient amounts to effect nuclear TOC1 turnover. The original report of endogenous ZTL cellular localization indicated a strictly cytosolic presence ^54^. A subsequent study relying on transgenic plants expressing ZTL HA-tagged protein reported detectable nuclear ZTL-HA ^55^. Further studies are planned to resolve the specific nuclear components responsible for the targeting of nuclear TOC1 to the proteasome.

## Materials and methods

### Plant materials

The *Arabidopsis thaliana* Columbia ecotype was used for all experiments. All *CCA1::LUC* lines used in this study were obtained by crossing transgenic plants with established *CCA1::LUC*^56^. To generate *NUA* co-suppression lines, the full-length *NUA* cDNA in pENTR/D-TOPO vector^31^ was subcloned into the gateway destination vector *gCsVMV-HA-1300*, under the control of *CsVMV* promoter, by performing LR recombination reaction (Invitrogen). The derived construct was transformed into *Agrobacterium tumefaciens* strain GV3101. WT *Arabidopsis* carrying *CCA1:LUC* reporter were transformed using the floral dip method ^57^. Homozygous transgenic lines with single insertion were used for further studies. To produce *nua-3* (SAIL_505_H11) and *nua-4* (WiscDsLox297300_17E) mutants expressing epitope-tagged proteins, *nua alleles* were crossed to *TOC1-YFP* minigene (TMG ^38^), *PRR3:PRR3-GFP* ^8^, *ELF4:ELF4-HA* ^58^, *35S:NUP136-GFP nup136-1*, and *35S:RAE1-GFP* ^35^.

To generate double mutants, *nua-3* was crossed with *siz1-2* (SALK_065397), *esd4-2* (SALK_032317), *cca1-1 lhy-20* ^59,60^ (kindly provided by the McClung lab), *toc1-101* ^61^, *prr9-1* (SALK_07551), *prr7-3* (SALK_030430), *prr5-1* (SALK_006280) and *prr3-1*..

### Plant growth conditions

For collecting tissues, surface-sterilized Arabidopsis seeds were plated on Murashige and Skoog (MS) media with 3% sucrose and 0.8% agar and incubated in darkness for 2-3 days at 4°C. The seedlings were grown in 12 h:12 h white-light cycles at 22°C in a growth chamber equipped with cool white fluorescent lamps until the tissues were collected. For normal plant growth, seedlings were transferred to soil and grown in a 16 h:8 h photoperiod (white light, 110 µmol m^−2^s^−1^) at 22°C.

### Plasmid construction

*TAP-TOC1* full length plasmid was generated in a previous study ^8^. To clone *TOC1* into pDEST-GADT7 deficient in its innate SV40 NLS site (pDEST-GADT7-ΔNLS), the *GAL4 AD* of pDEST-GADT7 was replaced by sequence without the SV40 NLS site via Hind III digestion and ligation. TOC full length sequence in pENTR/D-TOPO vector (Invitrogen) was moved to the pDEST-GADT7-ΔNLS by Gateway cloning (Invitrogen). Construction of plasmid pDEST22-NUA F1R2 containing *NUA* N-terminus (1-1248 aa) was described previously ^62^. pDEST22-NUA CT plasmid harboring NUA C-terminus fragments (1249-2093 aa) was kindly provided by Iris Meier (The Ohio State University, Columbus, OH, United States. Full length NUP136 coding sequence without STOP codon was cloned into pSITE-4NA mRFP binary vector (ABRC; CD3-1642) using LR recombination (Invitrogen). Full-length sequences of TOC1, PRR5, ELF4, and NUP136 were cloned into the pENTR/D-TOPO vector (Invitrogen). These entry clones were separately recombined into the Gateway destination vectors pEarleyGate101 (C-terminal YFP vector driven by the 35S promoter; from ABRC), pK7FWG2 (C-terminal GFP vector driven by the 35S promoter; gift from Dr. JC Jang), and pEarleyGate102 (C-terminal CFP vector driven by the 35S promoter; from ABRC) through LR recombination to generate TOC1-YFP, PRR5-YFP, ELF4-GFP, and NUP136-CFP constructs.

### Bioluminescence assays and hypocotyl length measurement

The seeds of *CCA1:LUC* and *PRR7:LUC* transgenic plants were sown on MS media containing 3% sucrose and grown in 12 h:12 h white-light cycles with 50 µmol m^−2^s^−1^ of white fluorescent light at 22°C. The 5-7-day-old seedlings were sprayed with 1 mM luciferin solution containing 0.01% Triton X-100 and imaged (Nightowl; Berthold Technologies, Bad Wildbad, Germany) at 22°C. Luminescence images were obtained every 2 h for seven days under continuous red LED light (30 µmol m^−2^s^−1^). Period length was estimated by FFT-NLLS analysis using BRASS3 (https://www.ed.ac.uk/profile/andrew-millar) and BioDare2 ^63^. For determination of hypocotyl length, seeds were sown on MS media without sucrose and placed in darkness for five days at 4°C for stratification. Plants were grown in continuous red LED light at various intensities ranging from 0.6 to 22 µmol m^−2^s^−1^. Hypocotyl lengths of five-day-old seedlings were measured using ImageJ.

### RNA extraction and quantitative real-time PCR

Total RNA was extracted from ground tissue using Trizol™ according to the manufacturer’s instruction (Thermo Fisher Scientific, 15596026), followed by DNase I treatment (Ambion, AM2224) for 30 min at 37°C. For cDNA synthesis, approximately 1 µg of RNA was incubated in a mixture containing Oligo(dT)^12–18^ primer and SuperScript™ III reverse transcriptase (Thermo Fisher Scientific, 18080093) for 60 min at 50°C, and treated with RNase H for 20 min at 37°C. Gene-specific primers and iQ™ SYBR Green Supermix (Bio-Rad, 1708882) were added to template cDNA and the readout fluorescence signal was analyzed by CFX Manager™ Software (Bio-Rad).

### Protein extraction and immunoblot analysis

Total protein extraction and immunoblot analysis were conducted as previously ^64^. Briefly, the ground tissues were homogenized with extraction buffer (50 mM Tris-Cl, pH 7.5, 150 mM NaCl, 0.5% Nonidet P-40, 1 mM EDTA, 1 mM dithiothreitol, 1 mM phenylmethylsulfonyl fluoride, 2 mM sodium fluoride, 2 mM sodium orthovanadate, 5 µg/ml leupeptin, 1 µg/ml aprotinin, 1 µg/ml pepstatin, 5 µg/ml antipain, 5 µg/ml chymostatin, 50 µM MG132, 50 µM MG115, and 50 µM ALLN). After centrifugation at 13,000 rpm for 10 min at 4 °C, the supernatant was combined with Urea/SDS loading buffer (40 mM Tris-Cl, pH 6.8, 8 M Urea, 5% SDS, 1 mM EDTA, 2% 2-mercaptoethanol) to be subjected to SDS-PAGE (acrylamide:bisacrylamide, 37.5:1). The separated proteins on the gel were transferred to PVDF membrane and probed with 1:5,000 anti-GFP (Abcam, ab6556), 1:1,000 anti-HA (Sigma, 3F10), 1:2,000 anti-myc (Sigma, M4439), 1:200 anti-NIP153 (BioLegend, MMS-102P), and 1:2,000 anti-WIT1 (Zhao et al., 2008). The anti-Histone H3 (Abcam, ab1791) and anti-ADK (gift from Dr. David Bisaro) were used as a loading control in 1:15,000 dilution.

### Subcellular fractionation and fluorescence signal-based measurement of nucleo-cytoplasmic ratio

The fractions of nuclear and cytoplasmic proteins were isolated using CELLYTPN1 CelLytic PN isolation/Extraction Kit (Sigma-Aldrich, CELLYTPN1-1KT). Homogenized tissues in nuclear isolation buffer (1X nuclear isolation buffer plus 1 mM dithiothreitol, 2 mM sodium fluoride, 2 mM sodium orthovanadate, 1 mM phenylmethylsulfonyl fluoride, 5 µg/ml leupeptin, 1 µg/ml aprotinin, 1 µg/ml pepstatin, 5 µg/ml antipain, 5 µg/ml chymostatin, 50 µM MG132, 50 µM MG115, and 50 µM ALLN), were aliquoted in 40 µl volume as a total fraction, and the rest of the mixture was filtered with four layers of Miracloth (Calbiochem, 475855). The filtrates were centrifuged at 1,000 x g for 10 min at 4°C, and the supernatant and pellet were separately transferred to fresh tubes. To obtain the cytoplasmic fraction, the supernatant was centrifuged at 13,000 rpm for 15 min at 4°C three times, and the resultant supernatant was mixed with the Urea/SDS loading buffer. For the nuclear fraction, the filtrated pellet was resuspended in nuclear isolation buffer additionally supplemented with 0.3% Triton X-100 and incubated on ice for 5 min at 4°C. After washing four times by repeating centrifugation at 6,800 rpm for 5 min at 4°C and resuspension in the same nuclear isolation buffer, the final pellet was resuspended in the above buffer by vigorous vortexing for 5 min and blended with Urea/SDS loading buffer for SDS-PAGE.

To analyze the nuclear/cytoplasmic ratio using confocal microscopy in Arabidopsis, 5-day-old seedlings were prepared on MS medium with 1% sucrose. Fluorescence images were acquired from 3-5 seedling root tips per line using a Nikon Eclipse C2plus confocal microscope with 488 nm excitation wavelength for GFP, 10% laser power, and a 40x oil immersion objective. Enlarged images were obtained using a 40x oil immersion objective plus 5x optical zoom. The detector gain was set to approximately 8.25 for comparable signal intensity. Signal intensities were quantified in ImageJ/FIJI, using the ROI tool, with ROIs of identical area for nucleus and cytoplasm. Mean of nuclear/cytoplasmic ratio was calculated from at least 70 cells per sample.

### Coimmunoprecipitation

For transient protein expression, *Agrobacterium tumefaciens GV3101* cells harboring *TAP-TOC1* were infiltrated into of 3–4-week-old *Nicotiana bethamiana* leaves. Tissues were collected three days after the infiltration and total protein extract were incubated with 20 ul of Human IgG-Agarose (Sigma, A6284) for 1 h at 4 °C. The resin was rinsed with cold extraction buffer four or five times and incubated with 1.5 ul of HRV3C protease (Thermo Scientific, 88947) for 1.5 h at 4°C with agitation to release immune complexes from the resin. After a brief spin down, the Urea/SDS loading buffer was added to the cleared supernatant.

#### Yeast two-hybrid and bimolecular fluorescence complementation assays

For Yeast two-hybrid assay, the bait/prey plasmids, pDEST-GBKT7/pDEST-GADT7 and pDEST32/pDEST22 were cotransformed to AH109 strain by the modified LiCl method^65^. The yeast cells were grown for 5-7 days at 30°C on minimal synthetic media (Fisher scientific, DF0919-15-3) supplemented with 2% glucose and 0.1-0.5 mM 3-AT but lacking adenine, histidine, leucine, and tryptophan, and the cell growth was evaluated. For BiFC assay, *Agrobacterium* GV3101 cells harboring either nYFP or cYFP clone along with H2B-RFP ^58^ were combined with the cells containing its counterpart clone or empty vector before infiltrated to *N. benthamiana* leaves. The fluorescence signal was observed three days after infiltration using confocal laser scanning microscopy.

#### Confocal microscopy

Four-to-five-week-old *N. benthamiana* leaves were infiltrated with Agrobacteria GV3101 and AGL-1 strains containing plasmids of interest. The OD600 of each culture was adjusted to 0.5 and combined in equal volumes for co-infiltrations, unless otherwise specified. Agrobacterium expressing gene-silencing repressor p19 were included in each infiltration. Sections from infiltrated leaves were mounted in ddH2O on microscope slides 72-96 h after inoculation. Arabidopsis nuclei expressing *NUP136:NUP136-GFP*, *WIP1:WIP1-GFP* and *35S:RAE1-GFP* were detected in root cells of 5-day old seedlings. Cells were imaged using a Leica Stellaris 5 confocal microscope fitted with a 40x water immersion objective (Leica Microsystems CMS, Germany). Image resolution was set to 512x512, 400 Hz scan speed and 4x zoom. CFP, GFP, YFP and mRFP were excited using lasers at 405 nm, 488 nm, 515 nm, and 560 nm, respectively, with emission detected at 456-510 nm (CFP), 490-550 nm (GFP), 518-545 nm (YFP) and 567-617 nm (mRFP). Z-stacks through nuclei were captured using the 40x objective set to 8x zoom for *N. benthamiana* and 4x for Arabidopsis. Images were acquired with LAS X software (Leica) and processed using ImageJ2 v2.16.0 and Icy v2.5.4.0.

### Statistical analysis

Student’s t-test and one-way ANOVA followed by Tukey’s HSD test were applied using R 3.6.3. Statistical significance was denoted by asterisks (Non-Significant (NS) *p* > 0.05, * *p* < 0.05, ** *p* < 0.01, *** *p* < 0.001).

## Data availability

The authors declare that all data supporting the findings of this study are available within the manuscript and its Supplementary Information files are available from the corresponding author upon request.

## Author contributions

YJ.K., B.M., J-W.Y., H.S., I.M. designed and performed the experiments, collected data, and analyzed the results. D.E.S. designed the project, some experiments, and wrote the manuscript. YS.L. contributed essential reagents.

## Acknowledgements

This work was supported by National Institutes of Health Grant R01GM093285 and R35GM136400 (D.E.S.) and by the National Research Foundation of Korea (NRF) grant funded by the Korea government (MSIT) (RS-2025-24533243, RS-2026-25497022), and by the New Faculty Startup Fund from Seoul National University (YS.L).

## Supplementary Materials

The PDF file includes: Supplementary Figures 1 to 14

