## Supplemental Figures 1-14 for "Nuclear basket localized proteasomes maintain circadian period through nuclear TOC1 proteolysis" (from another version of this preprint: version 1, 2026-05-23)

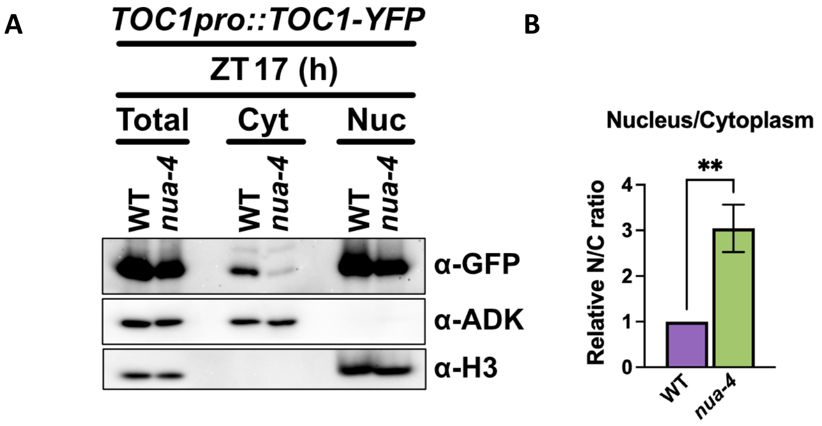


**Supplemental Figure 1. Subcellular distribution of TOC1 protein in the *nua-4* mutant.**

**(A)** Representative immunoblot showing TOC1 protein levels in total, cytoplasmic, and nuclear fractions from WT and *nua-4* plants using *TOC1pro::TOC1-YFP*. A representative result from three biological trials is shown.

**(B)** Quantification of TOC1 protein abundance in nuclear and cytoplasmic fractions from WT and *nua-4* plants. Data are presented as mean ± SD (n = 3). Statistical significance was determined by Student's t-test.

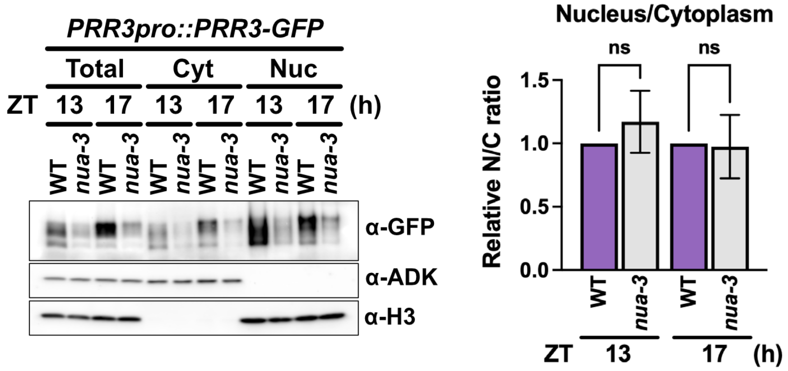


**Supplemental Figure 2. Nucleocytoplasmic distribution of PRR3 in *nua* mutants.**

Subcellular fractionation of *PRR3pro::PRR3-GFP* in WT and *nua-3* plants. Left panel shows representative immunoblots of total, cytoplasmic, and nuclear fractions. Right panel shows quantification of the nucleus-to-cytoplasm ratio. Data are presented as mean ± SD (n = 3). Statistical significance was determined by Student's t-test. A representative result from three biological trials is shown.

### 3D Projection Movie

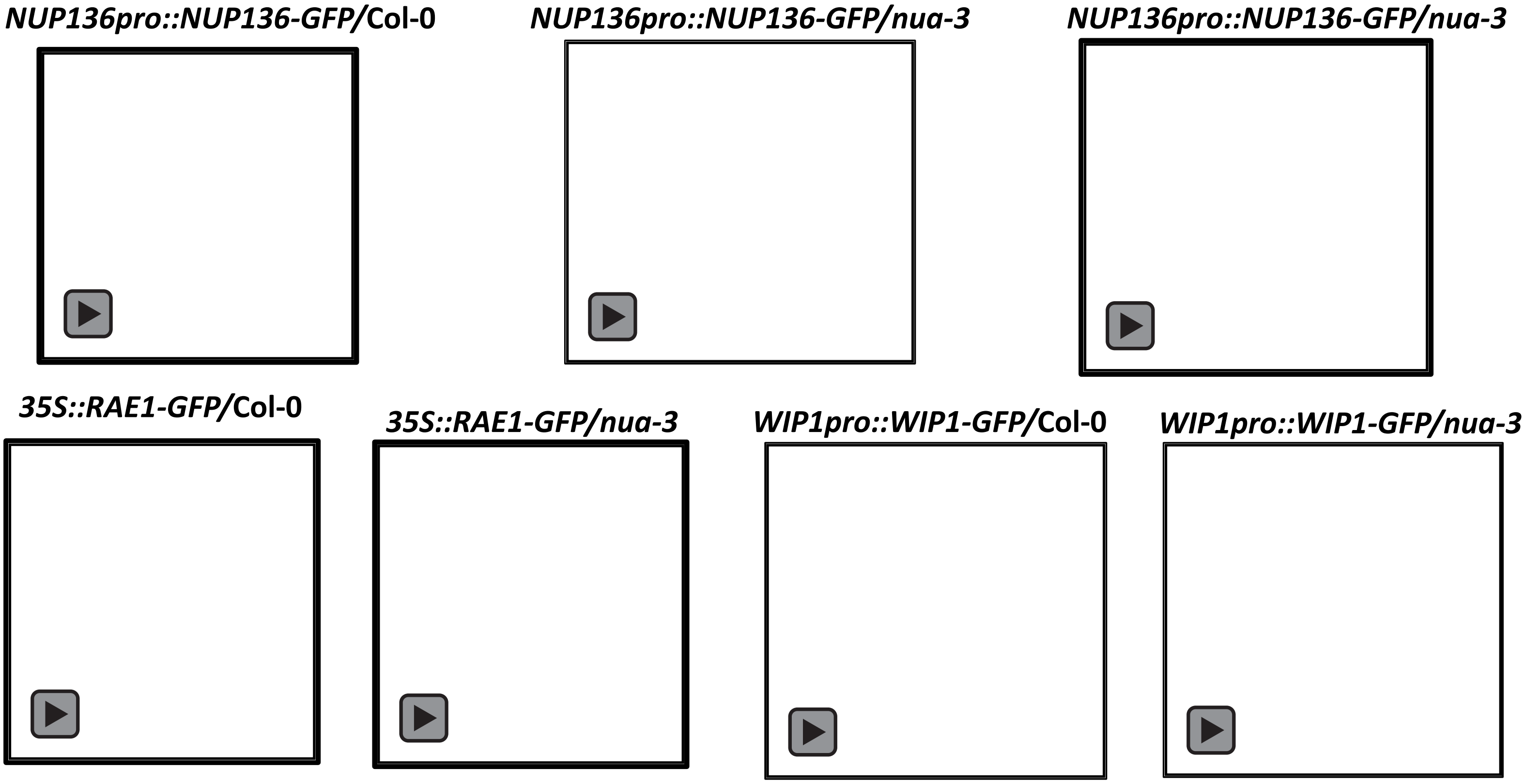


**Supplemental Figure 3. 3D-reconstructed images of nuclear envelope-localized GFP fusion proteins in *nua-3* mutant.**
3D-reconstructed images generated from serial confocal z-sections of root tips from 5-day-old seedlings. Top, *NUP136pro::NUP136-GFP* in Col-0 and *nua-3*. Bottom, *35S::RAE1-GFP* and *WIP1pro::WIP1-GFP* in Col-0 and *nua-3*.

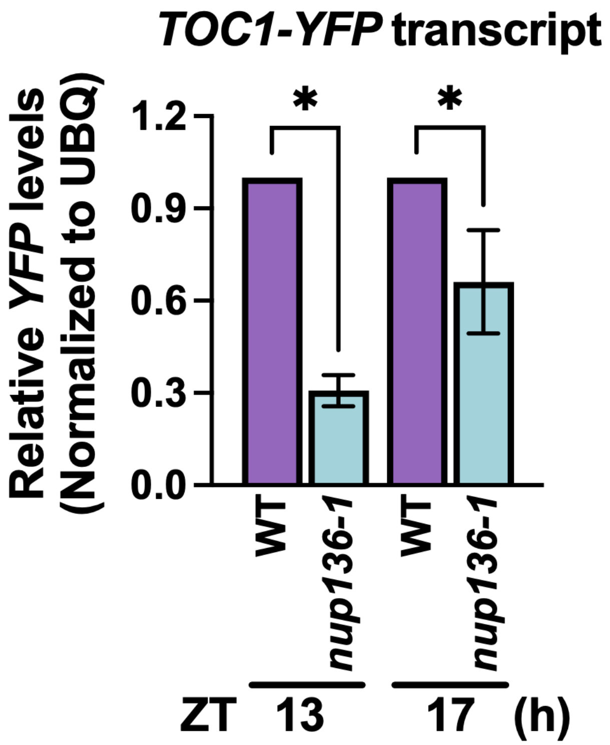


**Supplemental Figure 4. TOC1-YFP transcript levels in WT and *nup136-1* plants.** RT-qPCR analysis of *TOC1-YFP* transcript levels in WT and *nup136-1* plants. Samples were harvested at the indicated zeitgeber times (ZT). Data are presented as mean ± SD (n = 3).

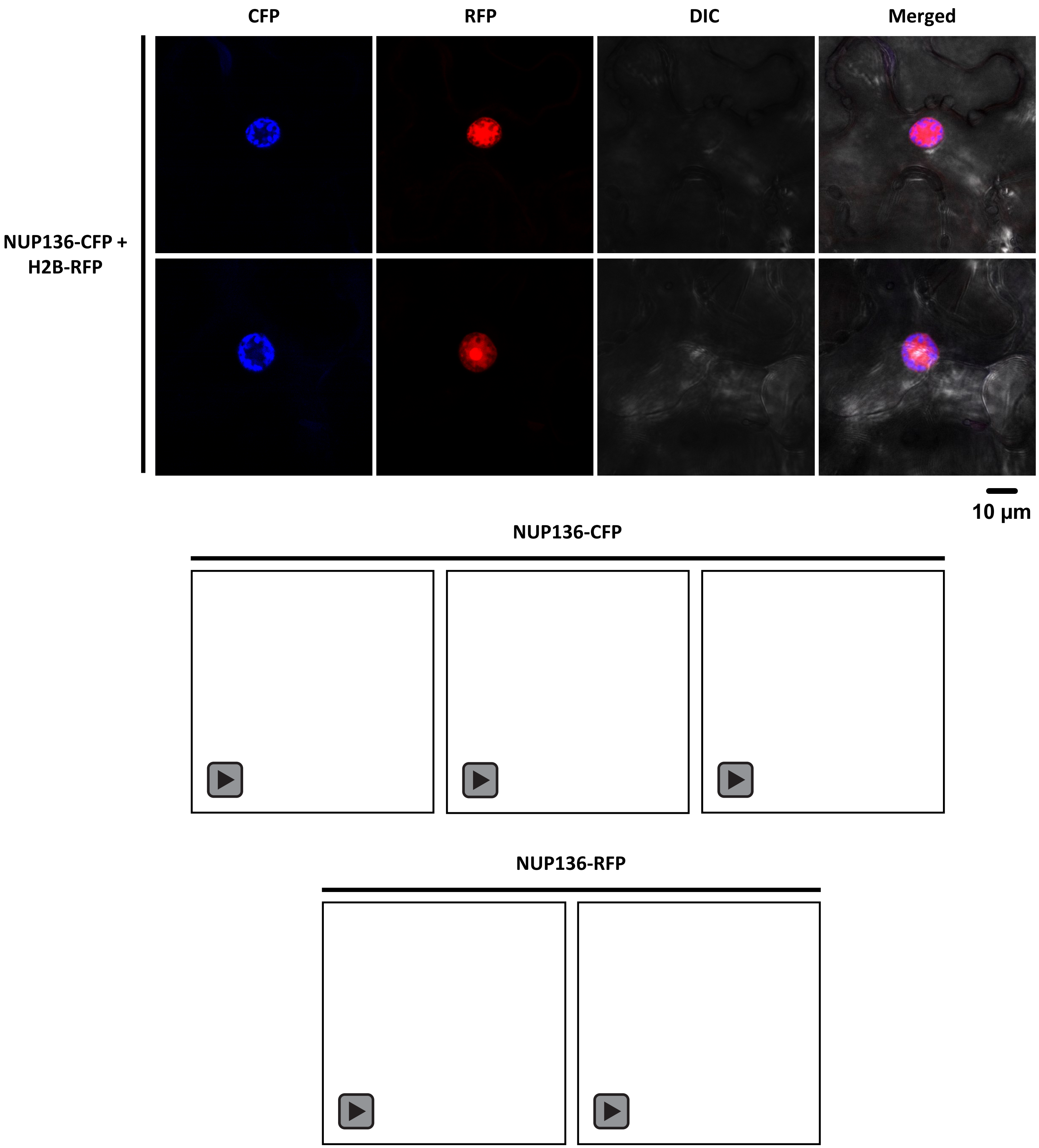


**Supplemental Figure 5. NUP136 nuclear localization**

**(A)** Nuclear localization of NUP136-CFP confirmed through transient co-expression of H2B-RFP in N. benthamiana leaf epidermal cells.

**(B)** 3D-reconstructed images of NUP136-CFP and NUP136-RFP showing nuclear envelope-localized fusion proteins**.** 3D-images generated from serial confocal z-sections.

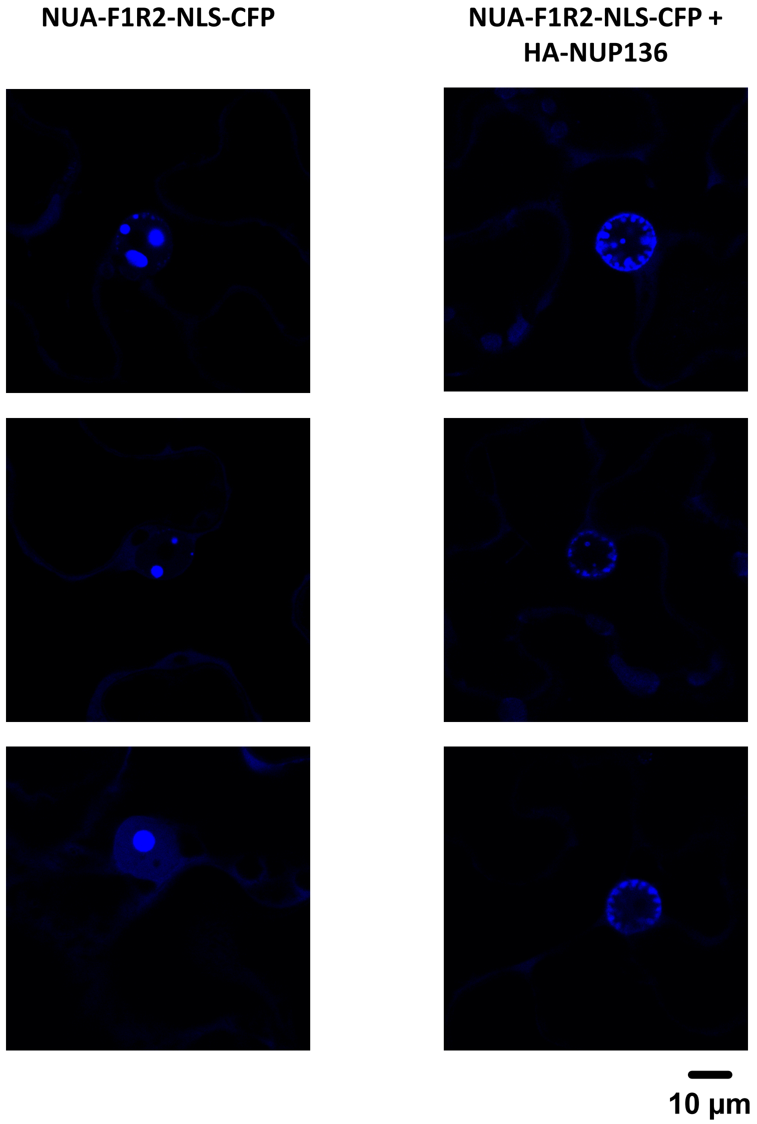


**Supplemental Figure 6. HA-NUP136 relocalizes NUA-F1R2-NLS-CFP to the nuclear periphery**
Confocal microscopy images of N. benthamiana leaf epidermal cells transiently expressing the indicated constructs at 3 days post-infiltration. Left, single expression of NUA F1R2-NLS-CFP. Right, co-expression of HA-NUP136 with NUA F1R2-NLS-CFP. Scale bars are indicated.

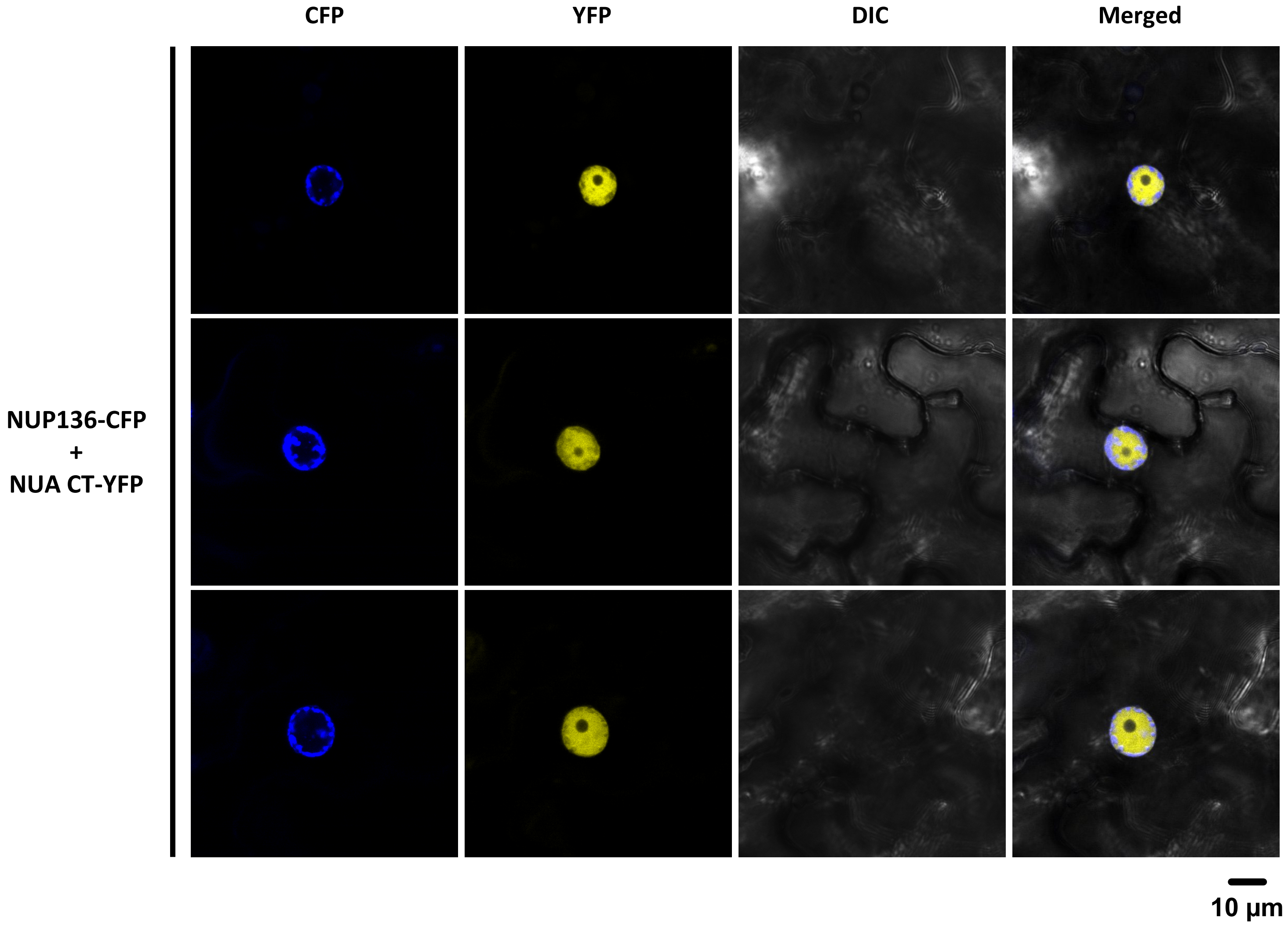


**Supplemental Figure 7. NUP136 does not relocalize NUA-CT to the nuclear periphery.**
Additional confocal microscopy images of *N. benthamiana* leaf epidermal cells transiently expressing the indicated constructs at 3 days post-infiltration. Scale bars are indicated.

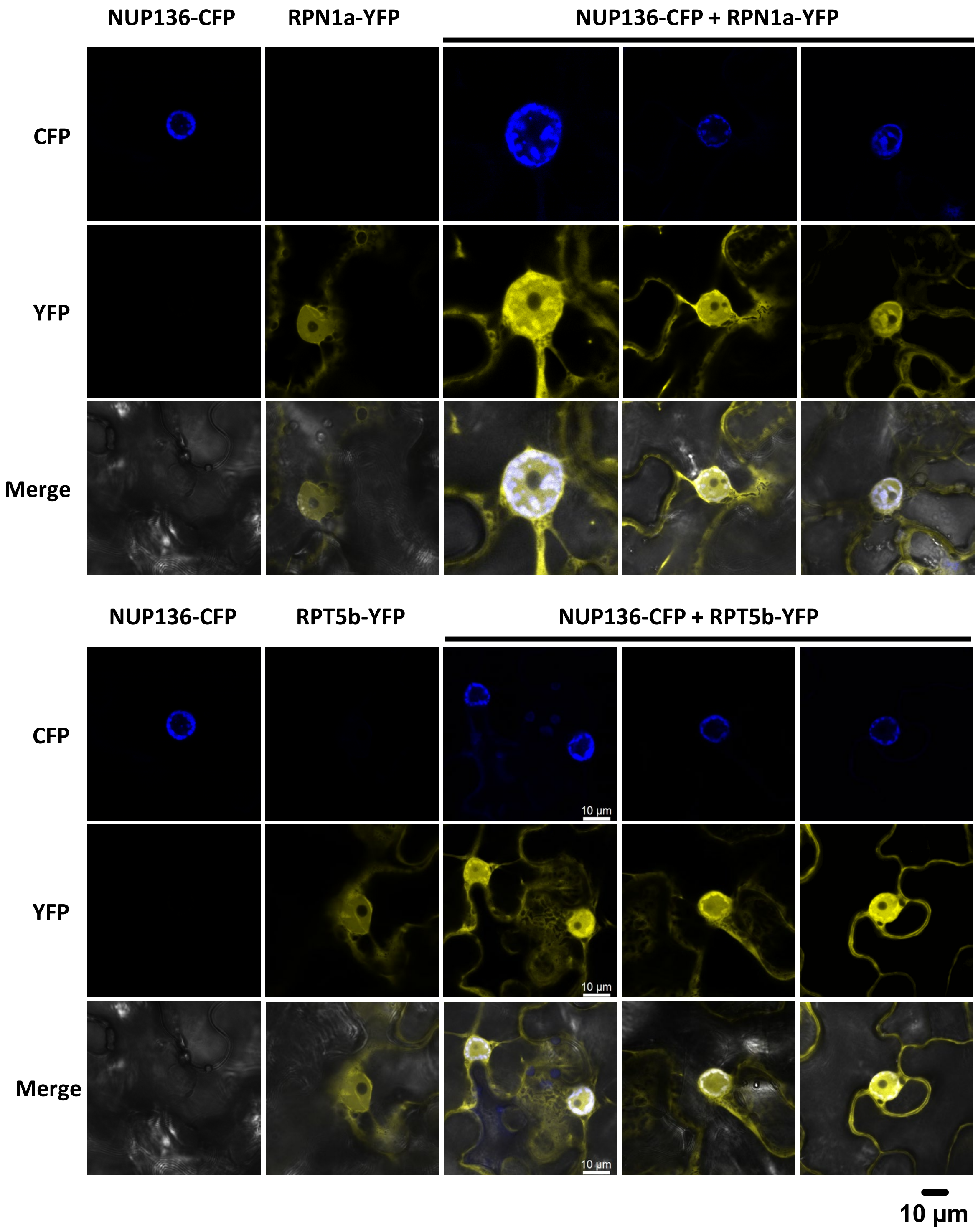


**Supplemental Figure 8. Additional colocalization images of NUP136 with proteasome subunits in *N. benthamiana* leaves.** Confocal microscopy images of *N. benthamiana* leaf epidermal cells transiently expressing the indicated constructs at 3 days post-infiltration. NUP136-CFP, RPN1A-YFP and RPT5B-YFP are shown alone and co-expressed. Scale bars are indicated.

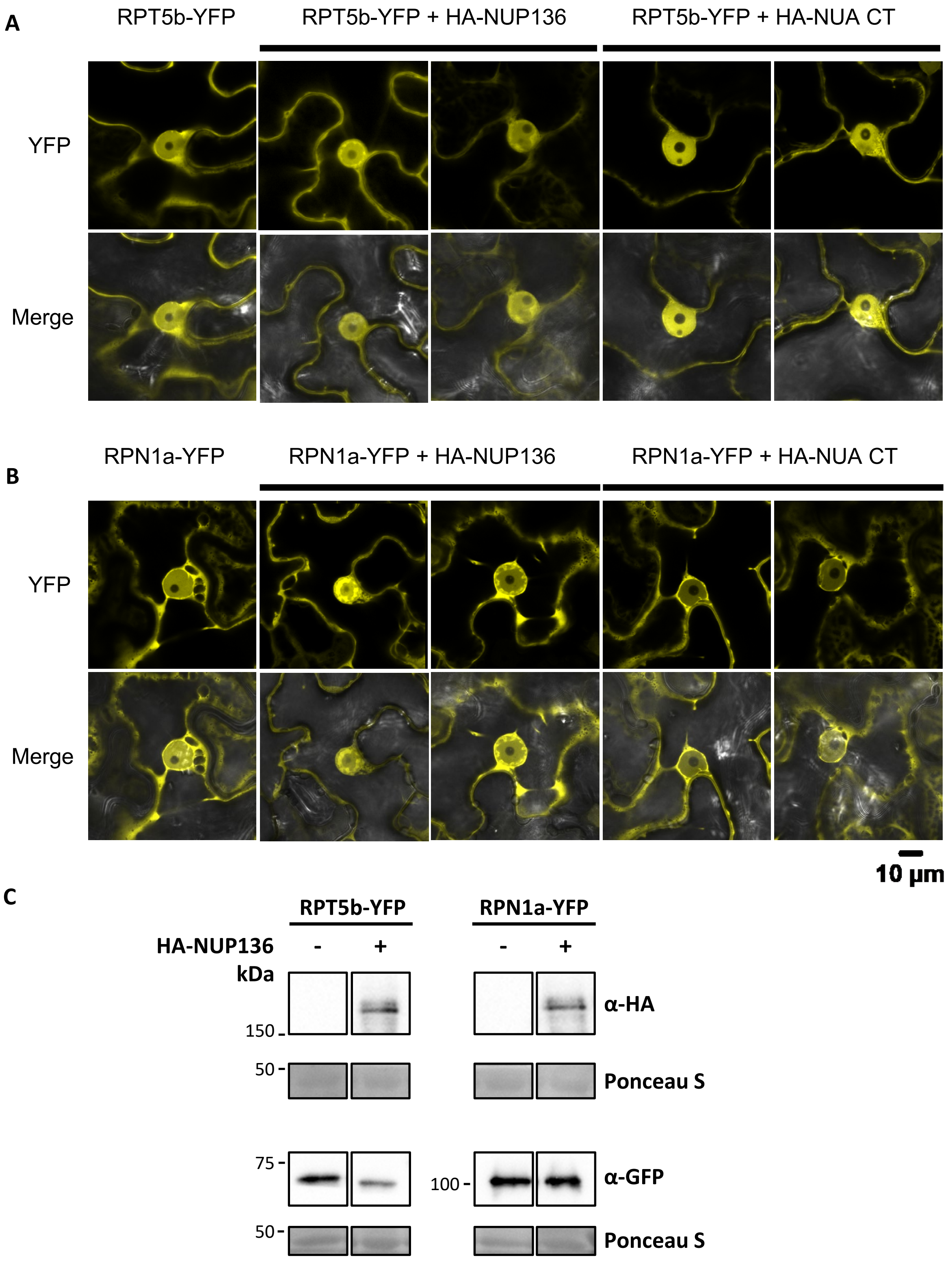


**Supplemental Figure 9. HA-NUP136 recruits proteasome components RPT5b and RPN1a to the inner nuclear periphery**

**(A)** RPT5b-YFP co-expressed with HA-NUP136 or HA-NUA-CT (negative control)

**(B)** RPN1a-YFP co-expressed with HA-NUP136 or HA-NUA-CT (negative control)

**(C)** Confirmation of HA-NUP136 and RPT5b/RPN1a-YFP expression in *N. benthamiana.* Immunoblots were probed with anti-HA (HA-NUP136) and anti-GFP (RPT5b/RPN1a-YFP). Ponceau S staining was used as a loading control.

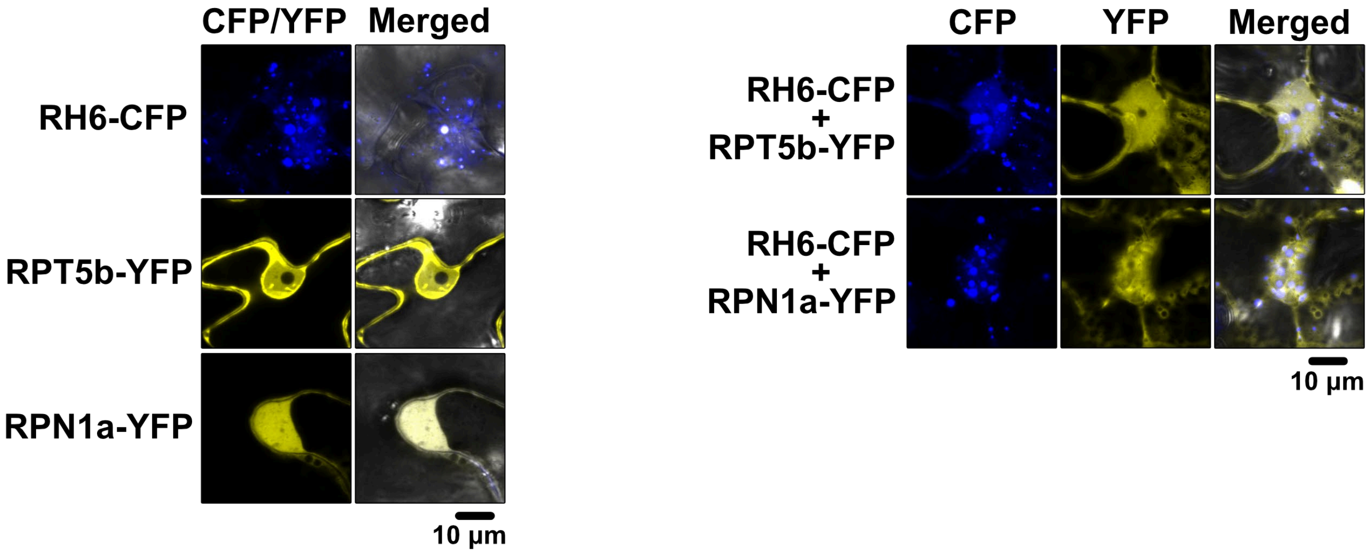


**Supplemental Figure 10. Colocalization analysis of RH6-CFP with RPT5B-YFP and RPN1A-YFP in *N. benthamiana* leaves.**

Confocal microscopy images of *N. benthamiana* leaf epidermal cells transiently expressing the indicated constructs at 3 days post-infiltration. RH6-CFP, a Dicer-associated protein that forms foci presumably distinct from those of NUP136, was included as a specificity control. Left, single expression of RH6-CFP, RPT5B-YFP, and RPN1A-YFP. Right, co-expression of RH6-CFP with RPT5B-YFP or RPN1A-YFP. CFP, YFP, and merged channels are shown. Scale bars are indicated.

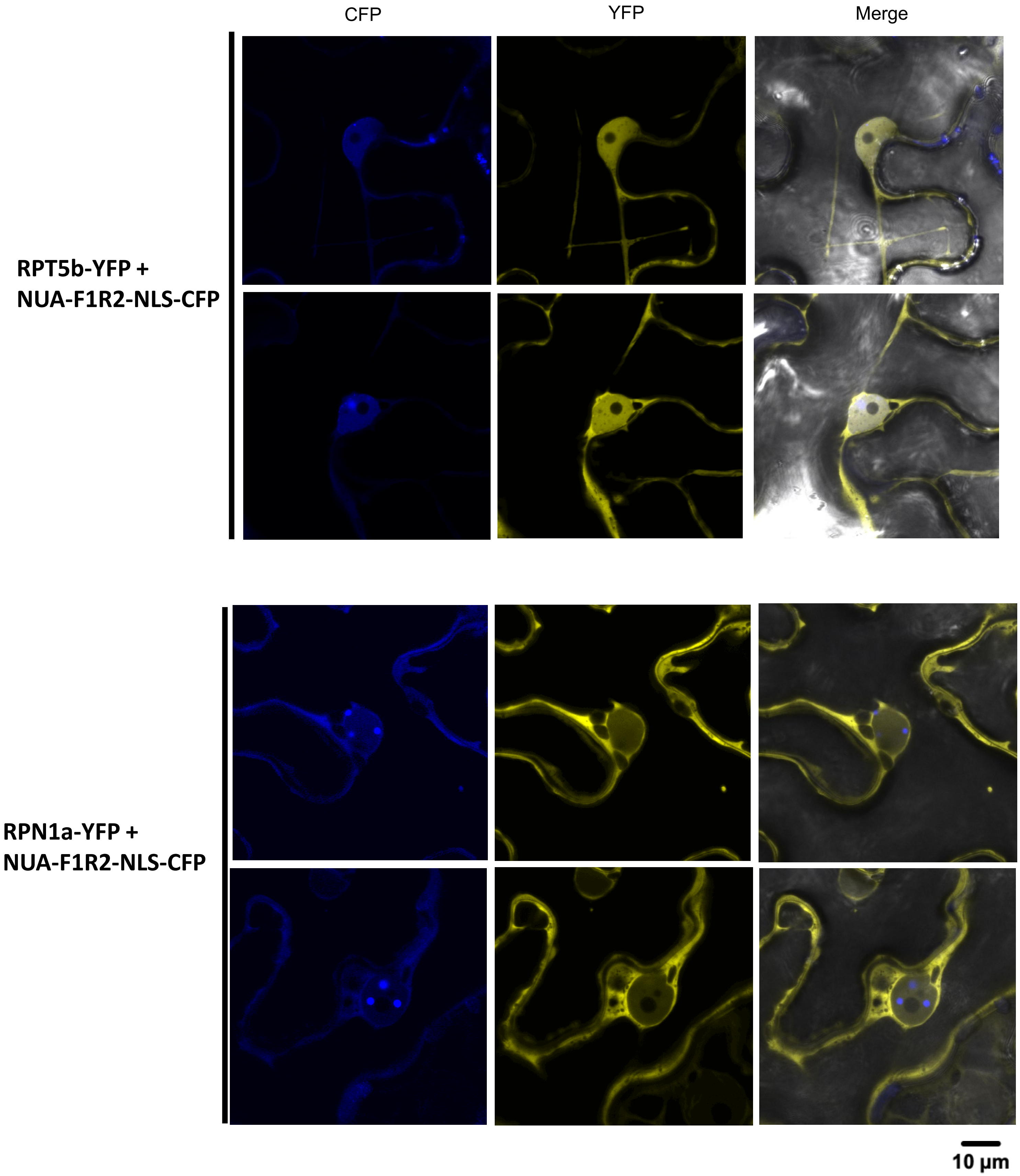


**Supplemental Figure 11. NUA N-terminus does not colocalize with RPT5B-YFP and RPN1A-YFP in *N. benthamiana* leaves.**

Confocal microscopy images of *N. benthamiana* leaf epidermal cells transiently expressing the indicated constructs at 3 days post-infiltration. Above, co-expression of RPN1a-YFP and NUA-F1R2-NLS-CFP. Below, co-expression of RPT5B-YFP with NUA-F1R2-NLS-CFP. CFP, YFP, and merged channels are shown. Scale bars are indicated.

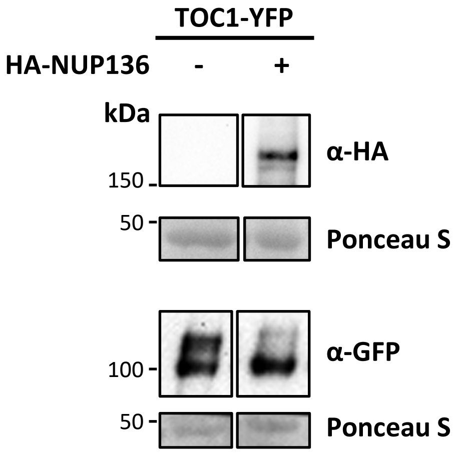


**Supplemental Figure 12. Confirmation of HA-NUP136 and TOC1-YFP expression in *N. benthamiana.*** Immunoblots were probed with anti-HA (HA-NUP136) and anti-GFP (TOC1-YFP). Ponceau S staining was used as a loading control.

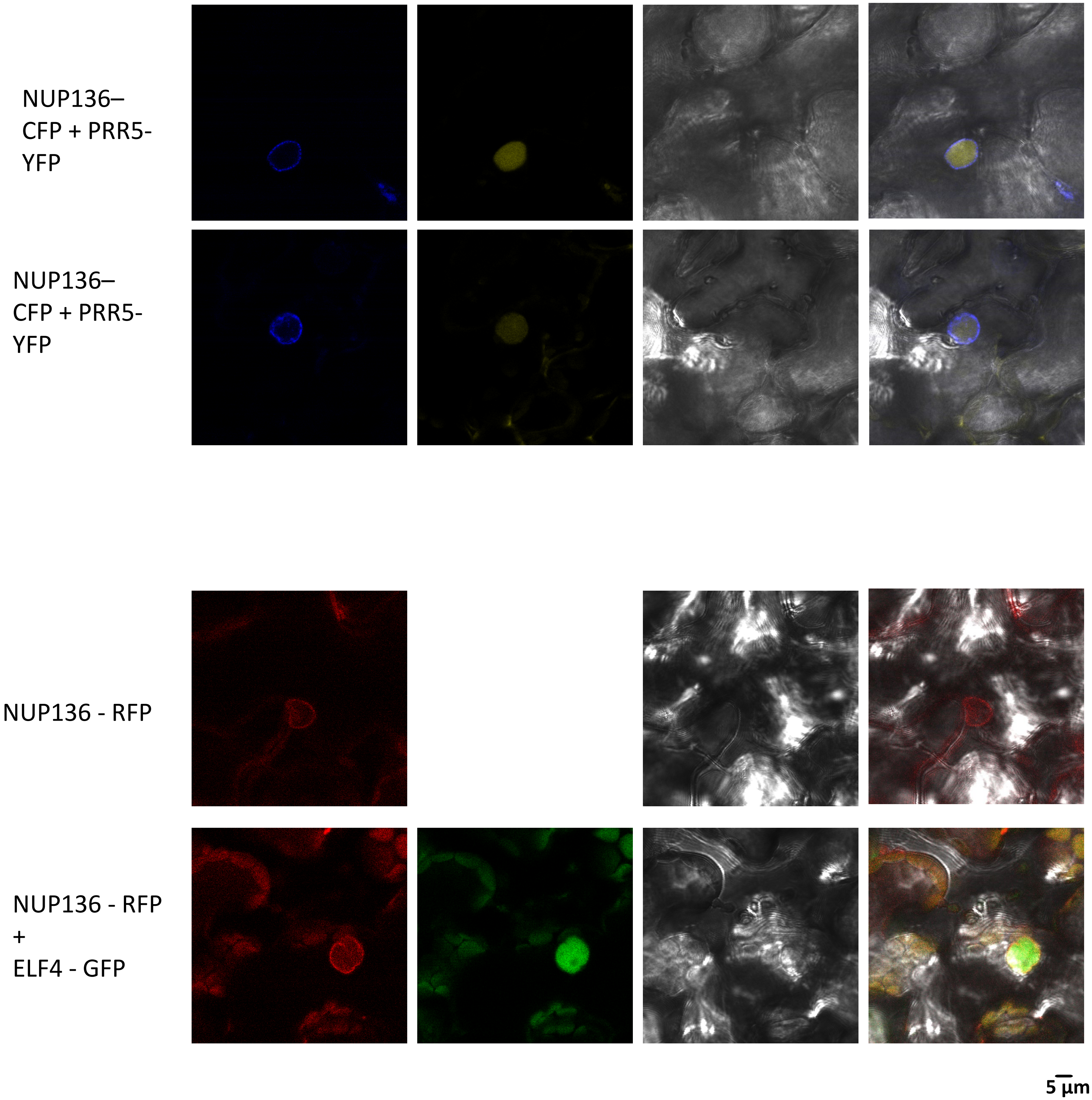


**Supplemental Figure 13**. **NUP136 does not alter PRR5 or ELF4 localization.**
Confocal microscopy images of N. benthamiana leaf epidermal cells transiently expressing the indicated constructs at 3 days post-infiltration. Scale bars are indicated.

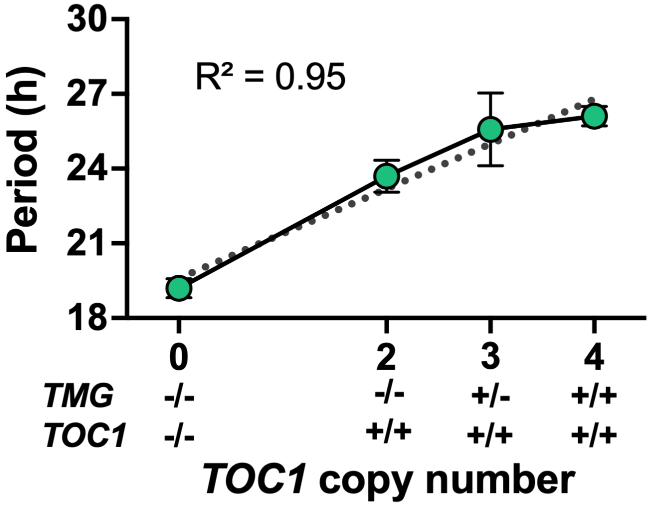


**Supplemental Figure 14. Relationship between *TOC1* dosage and circadian period.** Circadian period length plotted as a function of *TOC1* copy number in *TOC1pro::TOC1-YFP* lines. *TOC1pro::TOC1-YFP toc1-101* was outcrossed with Col-0, and period lengths were estimated in progeny lines harboring the indicated *TOC1* copy numbers. Data are presented as mean ± SD (n = 3).
